# Reproductive skew, fitness costs, and winner-loser effects in social-dominance evolution

**DOI:** 10.1101/2021.06.22.449392

**Authors:** Olof Leimar, Redouan Bshary

## Abstract

Social hierarchies can increase reproductive skew in group-living animals. Using game theory we investigate how the opportunity for differently ranked individuals to acquire resources influences reproductive skew, costs of hierarchy formation, and winner and loser effects. Individuals adjust their aggressive and submissive behaviour through reinforcement learning. The learning is based on perceived rewards and penalties, which depend on relative fighting ability. From individual-based simulations we determine evolutionary equilibria of traits that control an individual’s learning. We examine situations that differ in the extent of monopolisation of contested resources by dominants and in the amounts of uncontested resources that are distributed independently of rank. With costly fighting, we find that stable dominance hierarchies form, such that reproductive skew mirrors the distribution of resources over ranks. Individuals pay substantial costs of interacting, in particular in high-skew situations, with the highest costs paid by intermediately ranked individuals. For cases where dominants monopolise contested resources there are notable winner and loser effects, with winner effects for high ranks and very pronounced loser effects for lower ranks. The effects are instead weak when acquired resources increase linearly with rank. We compare our results on contest costs and winner-loser effects with field and experimental observations.

## Introduction

Social hierarchies often influence the distribution of reproductive success in group-living animals, with a skew towards higher success for dominant individuals (Ellis 1995; Clutton-Brock 1998; Clutton-Brock and Huchard 2013). The mating systems where dominance interactions can allocate reproductive success might extend beyond those of a group of individuals equally utilising an area, to also include systems with a spatial structure, such as leks and some forms of territoriality. To gain a broader perspective on empirical studies of such systems, and to inspire further investigation, it is of interest to derive theoretical predictions about the relation between an individual’s dominance rank and its reproductive success, as well as about the fitness costs of acquiring and maintaining a certain rank. Up to now there are, however, few theoretical models dealing with the distribution of reproductive success in social hierarchies, or with the costs of dominance interactions. Here we use an evolutionary game-theory model of hierarchy formation to examine reproductive skew, fitness costs, and winner and loser effects in social dominance.

Our model uses learning about differences in fighting ability as a behavioural mechanism that can give rise to within-sex dominance hierarchies, through pairwise interactions with aggressive and submissive behaviours. Learning in the model is actor-critic learning, which is a form of reinforcement learning (Sutton and Barto 2018). Individuals have genetically determined traits that function as parameters for the learning mechanism and can evolve to adapt learning to different situations. This approach has been used to study social dominance (McNamara and Leimar 2020; Leimar 2021) and here we extend it to allow for different degrees of monopolisation of reproduction by high-ranking individuals. We examine life histories with an annual life cycle. Over a season there are several reproductive cycles, each providing an opportunity for dominant individuals to monopolise reproduction. The situations we study range from an extreme case where all reproduction goes to a top-ranked individual in a group, over varying reproductive opportunities for lower-ranked individuals, to those where most reproduction is independent of rank and only a smaller part is achieved through winning pairwise contests. Our analysis could also apply to situations with nearby territories, or display sites on a lek, that differ in how valuable they are for reproduction and that are allocated according to a dominance hierarchy. The model might represent groups of males with mating opportunities as the resource that is contested, or females with foraging opportunities or nesting sites as the resource. Individuals are unrelated in the model, so it could apply to the dispersing sex in species where one sex disperses and the other is philopatric, and to either sex if both sexes disperse.

A basic desideratum for the model is that dominance hierarchies form in such a way that reproductive skew is higher when dominants have greater opportunities to monopolise reproduction. To describe reproductive skew we use the recently developed multinomial index (Ross et al. 2020). We also examine the statistical relation between rank and reproductive success, which can be more informative than a skew index. We use a variant of so-called Elo rating to indicate an individual’s rank. This measure was originally used for the ranking of chess players (Elo 1978) and is now often used to measure rank in social hierarchies (Albers and de Vries 2001; Neumann et al. 2011).

We express the cost of dominance interactions as mortality from fighting damage accumulated during the season, which means that individuals who die early in the season lose part of their reproductive opportunities. An equivalent effect would be a loss of vigour or condition from damage, eliminating reproductive success for the remainder of the season, or perhaps being weakened and driven away. We explore how fighting costs relate to reproductive skew and also how they depend on rank. For instance, costs could be higher for low-, medium-, or for high-ranked individuals. Based on what is known from previous game-theory models of social dominance, as well as from the long-standing study of single, pairwise contests, one would predict that the life-history costs of fighting should be higher when a greater proportion of lifetime reproductive success depends on winning dominance interactions. It is less clear how costs should depend on rank; there is no previous evolutionary analysis of this question. Studies on stress physiology in relation to social rank have found that subordinates are more stressed in some and dominants in other species (Creel 2001; Goymann and Wingfield 2004; Creel et al. 2013).

We also study if learning over a mating season affects an individual’s tendency to win or lose an interaction with a new, matched but naive opponent, and if this varies with the individual’s rank at the end of the season. This is related to and gives a new perspective on winner and loser effects, which have been much investigated experimentally (Rutte et al. 2006).

In the following, we briefly describe our model, present a number of results from individual-based evolutionary simulations, and discuss the implications of our results for observations of reproductive skew in social hierarchies and for the costs of dominance interactions. We also discuss how our model could be changed to take into account such things as multi-year life histories and overlapping generations, as well as the possible effects of common interest between members of a group.

## The model

Our model here is an extension of a previous one (Leimar 2021), with a new implementation of how dominance interactions occur over the mating season and how fitness benefits (reproductive success) and costs (mortality) come about. In the previous model, interactions in a group consisted of a sequence of rounds, each with randomly selected opponents, and fitness effects were represented as increments to payoffs (benefits and costs) that were translated into reproduction at the end of interactions. In the current model, interactions are structured into multi-round contests that occur during one or several reproductive opportunities or cycles, which might better correspond to natural interactions. Fitness effects are given a concrete life-history representation, with benefits as acquired resources, such as mating opportunities, and costs as mortality from accumulated fighting damage. Figure 1 gives an overview of these aspects.

**Figure 1:**
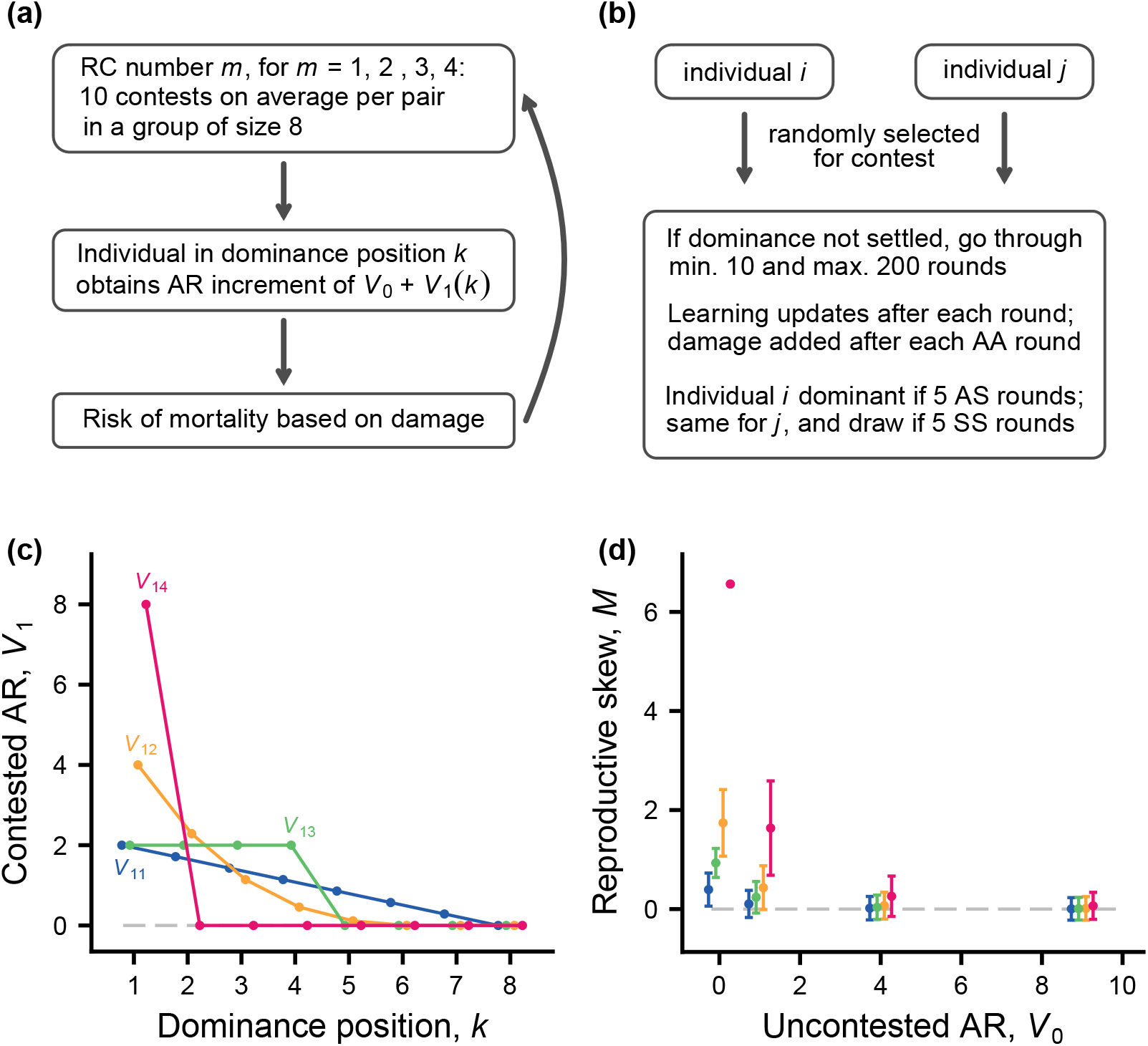
Elements of the model. During a mating season there are four reproductive cycles (RC) or reproductive opportunities, each starting with a sequence of contests, followed by increments in acquired resources (AR) and mortality risk, as shown in panel (a). The resource increments *V*_0_ and *V*_1_(*k*) contribute to reproductive success (RS). Panel (b) summarises a contest for a randomly selected pair of group members. Panels (c) and (d) illustrate the underlying distributions of AR and RS, for a single RC. (c) Increments *V*_1_(*k*), from contested resources, as functions of the dominance position *k*, where *k* = 1 is top-ranked. The curves *V*_11_, *V*_12_, *V*_13_, *V*_14_ (colour coded) show the different shapes of *V*_1_(*k*) used in simulations. For each curve, the mean per capita AR is 1. (d) Mean (± SD) of the multinomial reproductive skew index *M*, computed over 10 000 replicates of a group of size 8 that produces a total of 16 offspring (mean RS of 2 per group member), with AR given by *V*_0_ + *V*_1_(*k*). The skew values are shown as functions of the uncontested AR increment *V*_0_, for different shapes of *V*_1_(*k*), colour coded as in (c).

The season is divided into a number of reproductive cycles (Fig. 1a) in which group members participate. In a cycle, individuals meet in pairwise contests over dominance, with several contests per group member. The idea is that there should be good opportunities for group members to form a dominance hierarchy. For instance, a pair with similar fighting abilities can have several contests, potentially settling which of them dominates the other. A contest (Fig. 1b) can be thought of as an opportunity for dominance interaction; if dominance is already settled, there is no interaction. If there is an interaction, the model assumes a minimum and maximum number of rounds, to ensure that group members have experience of interacting with each other. A contest ends if there is a specified number of successive rounds with either a clear direction, so that one individual is aggressive and the other submits, which then indicates dominance, or a specified number of rounds where both submit, which indicates a draw. This aspect of the model is inspired by how dominance is often scored in experiments on hierarchy formation. The sequence of contests can produce a linear hierarchy, but it is also possible that there are cycles, or that some dominance relations remain undetermined, for instance if some group members avoid being aggressive towards each other, or if some continue fighting.

Following the contests, the resources (e.g., mating opportunities) are distributed according to rank (Fig. 1c). If some, or even all, ranks are undetermined at this stage, those ranks are randomly assigned (so if all individuals keep fighting, refusing to submit, resources are randomly acquired, both within and between cycles). We investigate four distributions of acquired resources over the ranks (Fig. 1c). They differ in how strongly the top ranks in a hierarchy monopolise resources, and are defined so that the mean acquired resource per individual is 1. The model also allows for uncontested resources, which are distributed to all (surviving) group members, irrespective of contest outcomes. The amount of reproductive skew that would results from these assumptions about acquired resources, for a hypothetical case where there is a single reproductive cycle with a linear dominance hierarchy, is shown in Fig. 1d.

The probability of survival from one reproductive cycle to the next depends on an individual’s accumulated damage at that point in time. Each round of fighting adds to damage, in a way that depends of the relative fighting abilities of the interacting individuals. When a new reproductive cycle starts (Fig. 1a), individuals are assumed to retain their previous learning, but the contests over dominance start as if from scratch. In many cases this results in the re-forming of a previous hierarchy with little or no fighting, but in some cases (e.g., when individuals have died) there can be contests with additional fighting. Finally, the reproductive success of each group member is allocated in proportion to its accumulated resources, for instance its mating success, over the entire season, irrespective of whether the individual survived the entire season.

For the Elo rating we let each individual *i* start with a rating of zero, *E_i_* = 0, and update ratings after an interaction resulting in dominance by *i* over *j* (Fig. 1b) by increasing *E_i_* and decreasing *E_j_* by 2.5(1 — *P_ij_*), with *P_ij_* = 1/(1 + exp(−*E_i_* + *E_j_*)). The ratings are also updated after a contest ending in a draw. This is a version of the approach previously used to measure rank (Albers and de Vries 2001; Neumann et al. 2011).

Important concepts and notation for the model are summarised in Table 1. A detailed model description, including those aspects that are the same as in the previous model (Leimar 2021), are presented in Supplementary Information Online.

**Table 1:**
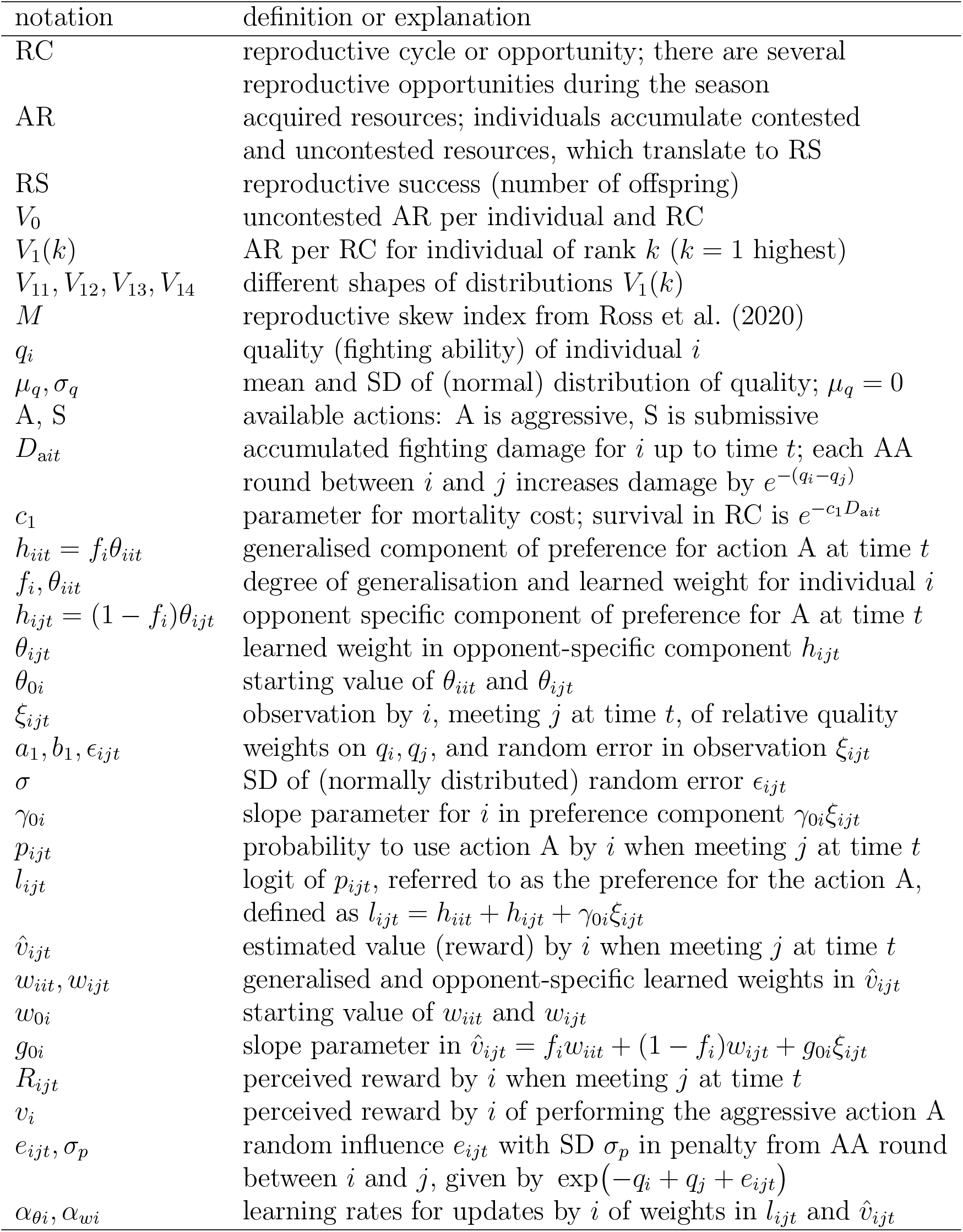
Definitions and notation for the model.

### Evolutionary simulations

As mentioned, individuals are assumed to have genetically determined traits. The evolution of the traits is studied in individual-based simulations. The traits for individual *i* are (Table 1): degree of generalisation, *f_i_*; preference and value learning rates, *α_θi_, α_wi_*; initial preference for action A, *θ*_0*i*_; initial estimated value, *w*_0*i*_; effect of observations on preference and value functions, *γ*_0*i*_, *g*_0*i*_; and perceived reward from performing A, *v_i_*.

In evolutionary simulations, each trait is determined by an unlinked diploid locus with additive alleles. Alleles mutate with a probability of 0.002 per generation, with normally distributed mutational increments. The standard deviation of mutational increments for each trait was adjusted to correspond to the range of values of the trait, to ensure that simulations could locate evolutionary equilibria.

A simulated population consisted of 500 groups of 8 individuals taking part in dominance interactions (either males or females), plus 8 individuals of the other sex, resulting in a total population size of *N* = 8000, which is the same as for simulations of the previous model (Leimar 2021). Each interacting individual was assigned a quality *q_i_*, independently drawn from a normal distribution with mean zero and standard deviation *σ_q_*.

Offspring for the next generation were formed by randomly selecting parents in a group for each of 16 offspring from that group, with probabilities proportional to an individual’s accumulated resources for the sex involved in interactions and uniformly for the other sex. The offspring were randomly dispersed over the groups in the next season, to eliminate any effects of relatedness in local groups. For each case reported in Table 2, simulations were performed over 5000 generations, repeated in sequence at least 100 times, to estimate mean and standard deviation of traits at an evolutionary equilibrium.

**Table 2:** Reproductive parameters, mean survival, and trait values (mean ± SD over 100 simulations, each over 5000 generations) for 16 different cases of individual-based evolutionary simulations of social dominance interactions.

| case | $V_0$ | $V_1$ | Surv. | $f_i$ | $\alpha_{\theta i}$ | $\alpha_{wi}$ |
| --- | --- | --- | --- | --- | --- | --- |
| 1 | 0.0 | $V_{11}$ | 0.64 | $0.035 \pm 0.008$ | $69.7 \pm 10.1$ | $0.036 \pm 0.005$ |
| 2 | 1.0 | $V_{11}$ | 0.71 | $0.045 \pm 0.011$ | $68.1 \pm 10.3$ | $0.047 \pm 0.008$ |
| 3 | 4.0 | $V_{11}$ | 0.81 | $0.068 \pm 0.017$ | $48.5 \pm 13.1$ | $0.076 \pm 0.016$ |
| 4 | 9.0 | $V_{11}$ | 0.87 | $0.108 \pm 0.027$ | $37.5 \pm 11.5$ | $0.123 \pm 0.031$ |
| 5 | 0.0 | $V_{12}$ | 0.51 | $0.073 \pm 0.015$ | $64.6 \pm 12.5$ | $0.035 \pm 0.006$ |
| 6 | 1.0 | $V_{12}$ | 0.62 | $0.131 \pm 0.012$ | $63.8 \pm 6.5$ | $0.039 \pm 0.006$ |
| 7 | 4.0 | $V_{12}$ | 0.74 | $0.191 \pm 0.019$ | $56.4 \pm 10.1$ | $0.054 \pm 0.010$ |
| 8 | 9.0 | $V_{12}$ | 0.82 | $0.372 \pm 0.018$ | $17.8 \pm 2.1$ | $0.045 \pm 0.012$ |
| 9 | 0.0 | $V_{13}$ | 0.61 | $0.025 \pm 0.005$ | $71.5 \pm 6.1$ | $0.032 \pm 0.005$ |
| 10 | 1.0 | $V_{13}$ | 0.68 | $0.032 \pm 0.008$ | $68.4 \pm 8.4$ | $0.039 \pm 0.007$ |
| 11 | 4.0 | $V_{13}$ | 0.78 | $0.054 \pm 0.013$ | $54.4 \pm 8.0$ | $0.058 \pm 0.011$ |
| 12 | 9.0 | $V_{13}$ | 0.85 | $0.076 \pm 0.018$ | $58.1 \pm 12.6$ | $0.098 \pm 0.026$ |
| 13 | 0.0 | $V_{14}$ | 0.39 | $0.249 \pm 0.021$ | $28.9 \pm 4.4$ | $0.028 \pm 0.009$ |
| 14 | 1.0 | $V_{14}$ | 0.56 | $0.484 \pm 0.018$ | $17.0 \pm 1.5$ | $0.016 \pm 0.004$ |
| 15 | 4.0 | $V_{14}$ | 0.68 | $0.572 \pm 0.016$ | $14.8 \pm 1.1$ | $0.014 \pm 0.003$ |
| 16 | 9.0 | $V_{14}$ | 0.77 | $0.613 \pm 0.018$ | $14.4 \pm 1.2$ | $0.017 \pm 0.003$ |

| case | $\theta_{0i}$ | $w_{0i}$ | $\gamma_{0i}$ | $g_{0i}$ | $v_i$ |
| --- | --- | --- | --- | --- | --- |
| 1 | $4.71 \pm 0.19$ | $-0.03 \pm 0.01$ | $1.23 \pm 0.18$ | $0.06 \pm 0.01$ | $0.86 \pm 0.01$ |
| 2 | $4.87 \pm 0.20$ | $-0.07 \pm 0.01$ | $1.84 \pm 0.23$ | $0.07 \pm 0.02$ | $0.76 \pm 0.02$ |
| 3 | $4.54 \pm 0.23$ | $-0.15 \pm 0.03$ | $2.79 \pm 0.34$ | $0.11 \pm 0.03$ | $0.56 \pm 0.04$ |
| 4 | $3.71 \pm 0.28$ | $-0.25 \pm 0.04$ | $2.96 \pm 0.35$ | $0.13 \pm 0.03$ | $0.42 \pm 0.05$ |
| 5 | $4.65 \pm 0.17$ | $-0.01 \pm 0.01$ | $0.64 \pm 0.20$ | $0.04 \pm 0.01$ | $0.95 \pm 0.01$ |
| 6 | $4.67 \pm 0.13$ | $-0.04 \pm 0.01$ | $0.95 \pm 0.18$ | $0.05 \pm 0.01$ | $0.89 \pm 0.01$ |
| 7 | $4.75 \pm 0.26$ | $-0.10 \pm 0.02$ | $1.78 \pm 0.29$ | $0.08 \pm 0.02$ | $0.76 \pm 0.02$ |
| 8 | $2.86 \pm 0.45$ | $-0.12 \pm 0.03$ | $1.66 \pm 0.37$ | $0.11 \pm 0.02$ | $0.68 \pm 0.02$ |
| 9 | $4.72 \pm 0.11$ | $-0.03 \pm 0.01$ | $0.92 \pm 0.18$ | $0.05 \pm 0.01$ | $0.89 \pm 0.01$ |
| 10 | $4.82 \pm 0.18$ | $-0.06 \pm 0.01$ | $1.57 \pm 0.22$ | $0.06 \pm 0.01$ | $0.81 \pm 0.01$ |
| 11 | $4.63 \pm 0.20$ | $-0.12 \pm 0.02$ | $2.41 \pm 0.30$ | $0.10 \pm 0.02$ | $0.64 \pm 0.03$ |
| 12 | $4.40 \pm 0.31$ | $-0.20 \pm 0.03$ | $3.27 \pm 0.52$ | $0.12 \pm 0.03$ | $0.49 \pm 0.05$ |
| 13 | $4.77 \pm 0.24$ | $0.00 \pm 0.02$ | $0.48 \pm 0.25$ | $0.05 \pm 0.02$ | $0.98 \pm 0.01$ |
| 14 | $3.17 \pm 0.24$ | $-0.05 \pm 0.02$ | $0.77 \pm 0.22$ | $0.07 \pm 0.02$ | $0.94 \pm 0.01$ |
| 15 | $0.76 \pm 0.52$ | $-0.06 \pm 0.03$ | $0.55 \pm 0.16$ | $0.08 \pm 0.02$ | $0.85 \pm 0.01$ |
| 16 | $-1.31 \pm 0.32$ | $-0.01 \pm 0.03$ | $0.73 \pm 0.17$ | $0.10 \pm 0.02$ | $0.70 \pm 0.02$ |

### Standard parameter values

The following ‘standard values’ of parameters (Table 1) were used: mortality cost from damage, *c*_1_ = 0.002; distribution of individual quality, *σ_q_* = 0.50; observations of relative quality, *a*_1_ = *b*_1_ = 0.707, *σ* = 0.50; perceived penalty variation, *σ_p_* = 0.25. For these parameter values, around 50% of the variation in the observations *ξ_ijt_* by individuals in each round is due to variation in relative fighting ability, *q_i_* — *q_j_*.

## Results

Using four distributions of resources over dominance ranks (*V*_1_(*k*), Fig. 1c) in combination with four values of uncontested resources (*V*_0_ = 0, 1, 4, 9), we analysed 16 cases of individual-based evolutionary simulations, summarised in Table 2. The course of interactions over the season is illustrated in Fig. 2, for the cases with *V*_0_ = 0. Time in the season is defined such that the reproductive cycles start at *t* = 0.00, 0.25, 0.50, 0.75. As can be seen, most of the divergence in Elo ratings occurs early in the first cycle (Fig. 2a), and this is also when most damage is incurred (Fig. 2b). The explanation is that there are more and longer fights early in the season. It should be noted that there is considerable variation between groups, depending on such things as the particular fighting abilities *q_i_* in a group, the timing of mortality, and randomness in contest outcomes (Fig. S1 gives examples).

**Figure 2:**
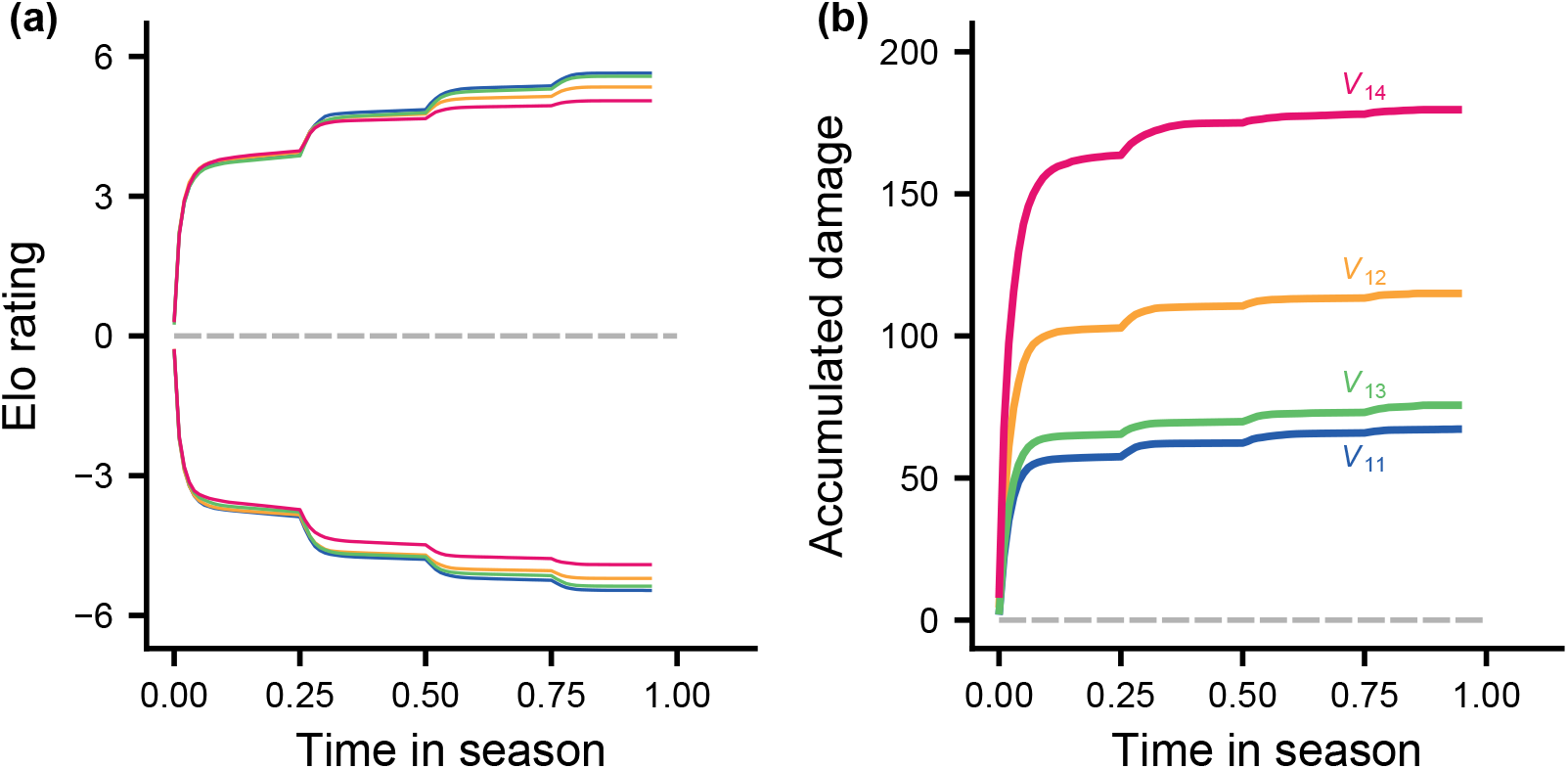
Average top- and bottom-ranked Elo ratings and average accumulated damage as functions of time in the season. The cases 1, 5, 9, and 13 in Table 2 (with *V*_0_ = 0) are shown, colour coded corresponding to the shapes of *V*_1_ in Fig. 1c. The learning parameters are given by the mean values in Table 2. For each case, 500 groups of 8 individuals were simulated and the Elo rating and accumulated damage of each individual as a function of time was computed (Fig. S1 shows examples of such curves). (a) Mean over groups of the Elo rating of the top (upper curves) and bottom (lower curves) ranked individuals in each group as functions of time in the season. (b) Mean over all groups and group members of the accumulated damage from fighting as functions of time in the season. Note that most damage accumulates in the first of the four reproductive cycles.

In presenting results, we show statistical model fits (non-linear regressions, including loess regressions), to ease comparison. Because there is considerable random variation in reproductive and damage outcomes between groups and individuals, we also show individual data points together with fitted curves in Supplementary Information Online.

Figure 3 gives an overview of the distribution of reproductive success and the cost of fighting for different cases (Figs. S2 and S3 show data points for panels (a) and (c)). The distributions of reproductive success for the different cases (Fig. 3a) have similar shapes as the underlying distributions of resources over dominance ranks (Fig. 1c). The reason is that dominance hierarchies remain fairly stable over the season, at least with regard to the rankings that matter for reproductive success. This also holds for the skew indices (Fig. 3b vs. 1d). The accumulated damage is highest for individuals of intermediate rank (Fig. 3c), and this is most pronounced for the case with the greatest opportunities for dominants to monopolise reproduction (*V*_14_). The mortality costs from damage are overall substantial (Fig. 3d), and are higher when a greater proportion of life-time reproduction can be acquired through social dominance.

**Figure 3:**
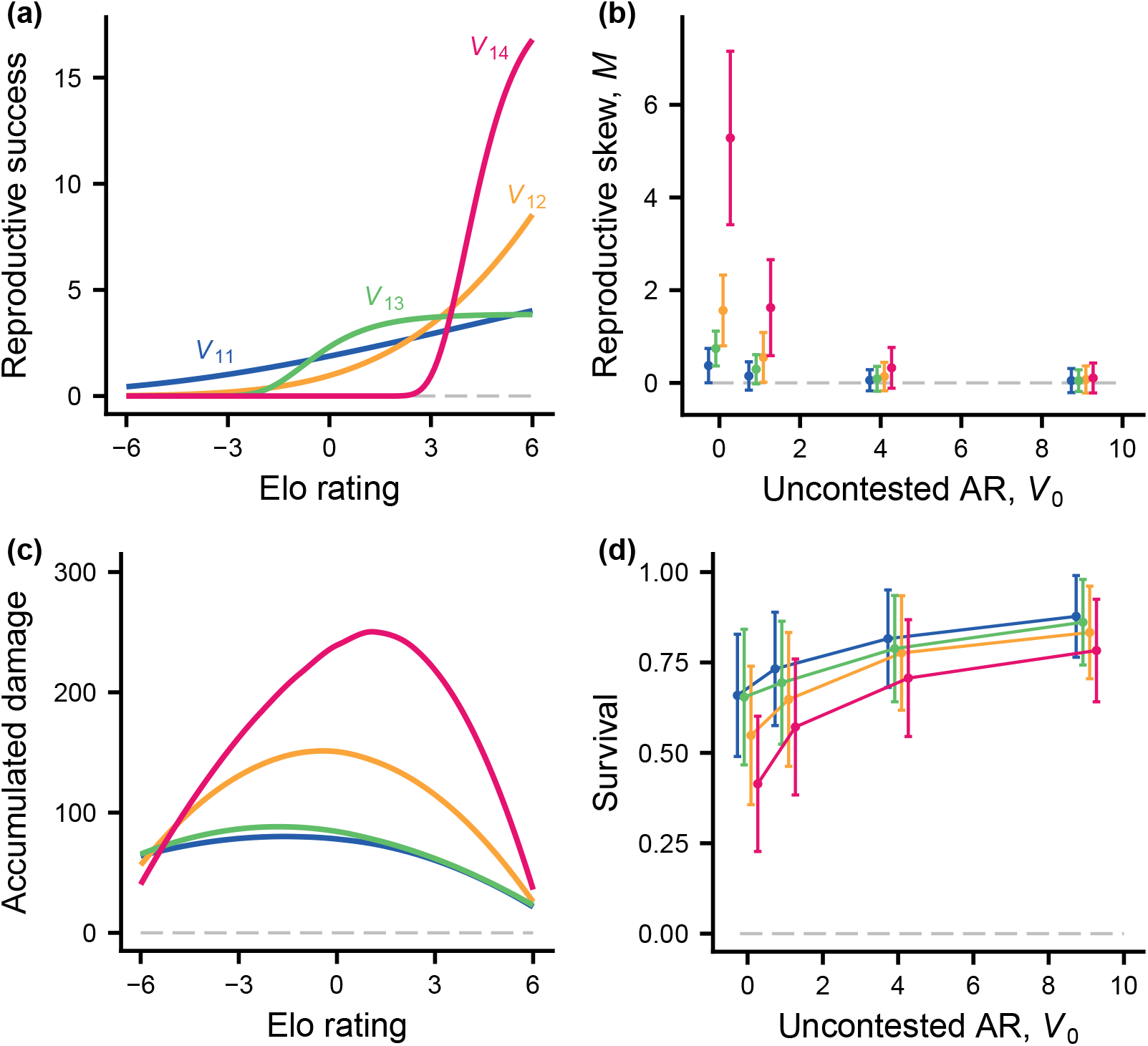
Distribution of RS (number of offspring; 16 per group), accumulated fighting damage, and survival, for different evolved learning parameters, with different distributions of contested and uncontested AR. Learning parameters are given by mean values in Table 2. For each case in Table 2, 500 groups of 8 were simulated. (a) Fitted number of offspring as a function of the Elo rating (which correlates with dominance position). The curves correspond to cases 1, 5, 9, and 13 in Table 2 (with *V*_0_ = 0) and are colour coded and labelled corresponding to the shapes of *V*_1_ in Fig. 1c. Fig. S2 shows simulated data together with fitted curves. (b) Mean (± SD) of the multinomial reproductive skew index *M*, computed from the distribution of RS in groups, for the cases in Table 2, with colour coding according the shape of *V*_1_ and the value of *V*_0_ along the x-axis (points are shifted left and right for clarity). (c) Fitted accumulated fighting damage over the season as a function of the Elo rating, for the cases in panel (a). Fig. S3 shows simulated data together with fitted curves. (d) Mean (± SD) survival in groups for the cases in Table 2, with colour coding according the shape of *V*_1_.

To examine winner and loser effects for different cases, we simulated experiments where group members who survived over the season met new, matched opponents in staged contests. The probability of winning (becoming dominant) and the damage for a group member in such contests can be influenced by learning from previous interactions, involving the effects of generalising from previous winning and losing. We investigated how these effects varied with the final Elo rating of a group member. As can be seen in Fig. 4a, winner and loser effects are strongest for the *V*_14_ case, followed by *V*_12_ and, with weaker effects, *V*_13_ and *V*_11_. Variation in accumulated damage shows a similar pattern (Fig. 4b). The explanation in terms of learning appears in Fig. 4c. Because of individual recognition, only the the generalised components of the action preference and the estimated value influence interactions with new opponents. The generalised component of the preference for action A is *h_iit_* = *f_i_θ_iit_*, where *f_i_* is the genetically determined degree of generalisation and *θ_iit_* is a learned weight (Table 1). This component will vary strongly with Elo rating if *f_i_* is large and *θ_iit_* shows notable variation, from negative to positive, with Elo rating. Effects of learning on *θ_iit_* will be larger when *f_i_* is larger, so the value of *f_i_* is driving the difference between the cases in Fig. 4c, and thus explains the results in panels (a) and (b) of the figure. The mean values of *f_i_* appear in Table 2 (cases 1, 5, 9, 13) and are in accordance with this explanation. Panels (a) and (c) of Fig. 4 also show that loser effects are stronger than winner effects, in the sense that the intersections with the horizontals Pr(win) = 0.5 and *h_iit_* = 0, respectively, occur for positive Elo ratings. Thus, on average group members tend to lose against matched opponents. The relation between fighting ability and Elo rating (Fig. 4d) shows that the final ratings are distributed approximately symmetrically around zero.

**Figure 4:**
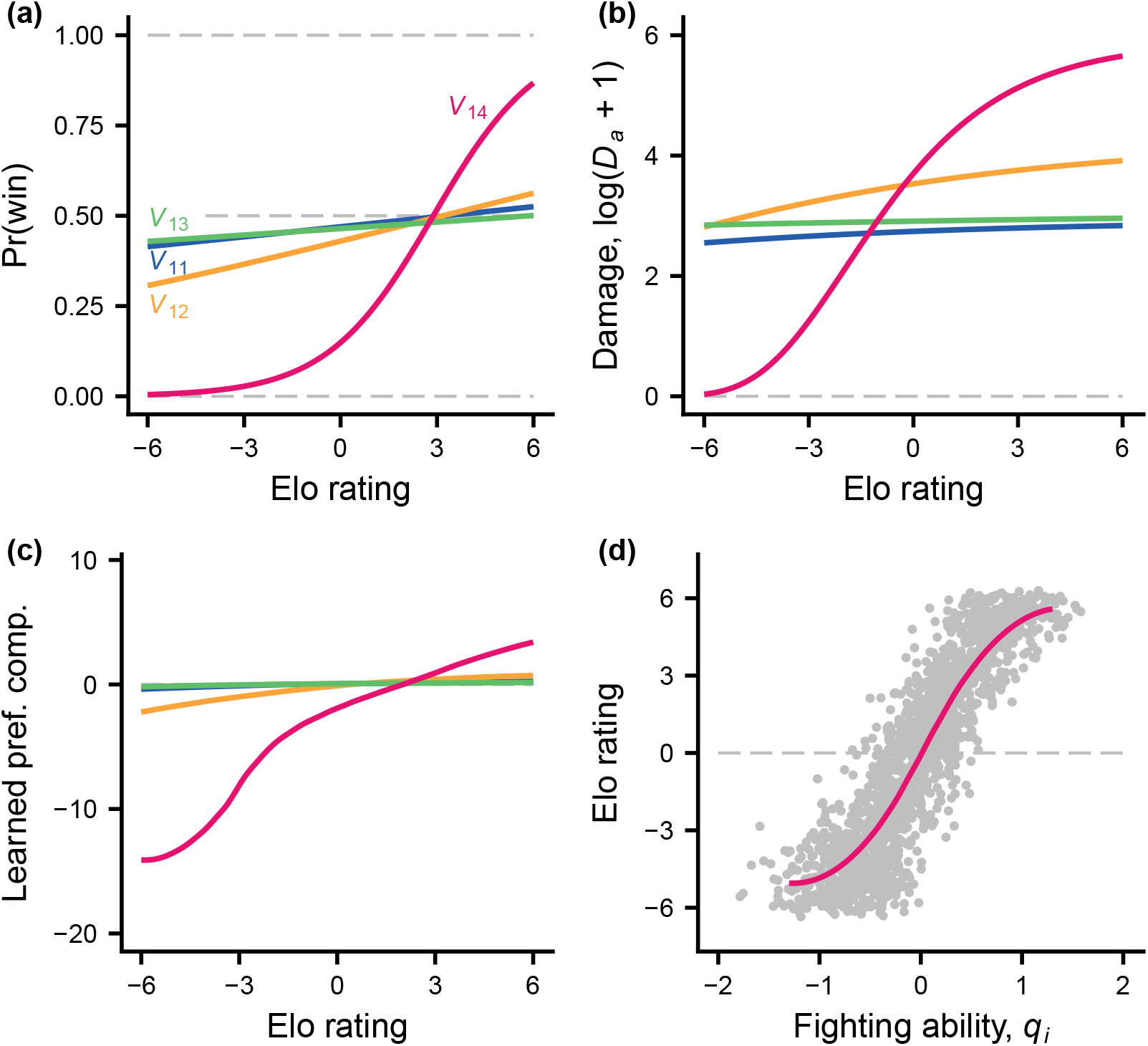
Illustration of hypothetical winner- and loser-effect experiments. Each group member that survived over the mating season had a staged interaction with a matched (equal fighting ability, *q_i_* = *q_j_*) new and naive opponent. A staged pair had up to 10 contests as described in Fig. 1b, ending when dominance was settled. The different cases (colour coded) are those in Fig. 2a, all having *V*_0_ = 0. For each case there were 500 simulated groups, including winner-loser experiments. (a) Fitted (logistic regression) probability of winning (becoming dominant) for a group member interacting with a matched, naive opponent, as a function of the group member’s Elo rating. (b) Fitted accumulated damage (logarithmic scale) from these contests. Fig. S4 shows simulated data together with fitted curves. (c) Fitted generalised preference component (*h_iit_* in Table 1) for group members at the start of staged interactions, as a function of Elo rating. Fig. S5 shows simulated data together with fitted curves. (d) Data and a loess fitted curve of the Elo rating as a function of fighting ability *q_i_* of group members, for case 13 in Table 2 (i.e., *V*_14_ in panels (a) to (c)).

The fitted curves in Figs. 3 and 4 are for cases with *V*_0_ = 0. With more of life-time reproduction coming from uncontested resources, the dependence of reproductive success on Elo rating is more shallow (Fig. 5a,b) and there is less fighting damage (Fig. 5c,d). In contrast, the degree of generalisation becomes higher for greater *V*_0_ (Table 2), resulting in stronger winner and loser effects (Fig. S6 and S7). Increasing *V*_0_ also resulted in lower values for the initial aggressiveness (*θ*_0*i*_ in Table 2).

**Figure 5:**
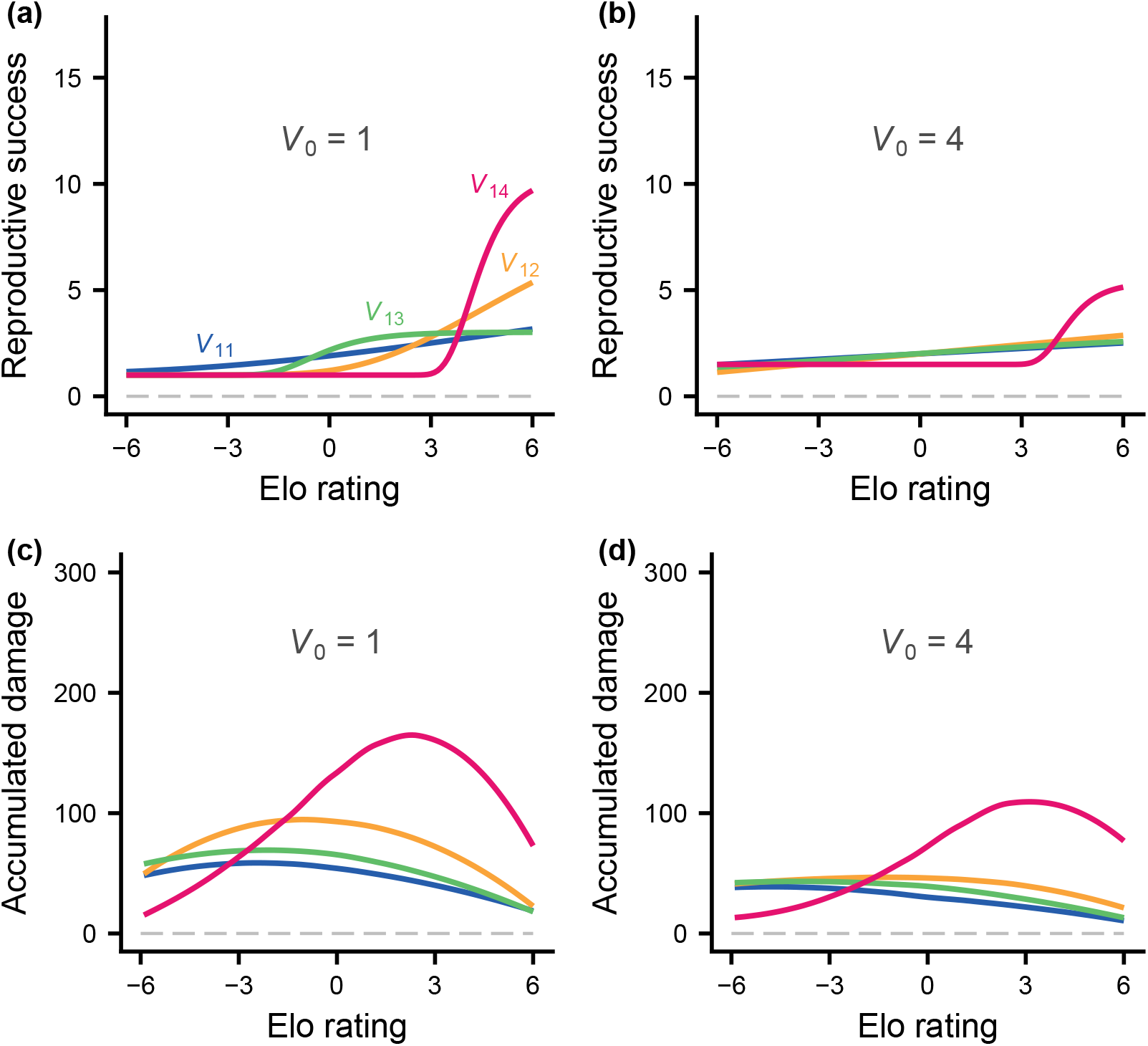
Fitted RS and accumulated damage over the season, as in Fig. 2a, c except that individuals also accumulate uncontested resources. Panels (a) and (c) show cases 2, 6, 10, and 14 in Table 2, with *V*_0_ = 1, and (b) and (d) show cases 3, 7, 11, and 15, with *V*_0_ = 4. For these values of *V*_0_, approximately 50% and 80% of a group’s total reproductive success derives from uncontested AR.

## Discussion

We found that the pattern of resource availability over the ranks of a hierarchy strongly influenced the evolution of learning mechanisms and, as a consequence, the costs of acquiring dominance. Because hierarchies remained relatively stable over the season, the distribution of reproductive success over ranks mirrored that of the resources (Fig. 3a vs. 1c). When dominants had greater opportunities to monopolise the resources needed for reproduction, there was greater skew (Fig. 3a, b). With greater skew, contests over dominance became more costly, in particular for individuals that had intermediate fighting abilities and were not among the top ranked (Fig. 3c, 5c, 5d). Such individuals have something to fight for, but face stiff competition. Winner and loser effects also depended on the pattern of resource availability and took the form of loser effects for lower-ranked individuals and winner effects for the top ranks (Fig. 4, S6, S7).

To appreciate these results, it is helpful to consider the reproductive success of all ranks of a hierarchy. A characteristic feature of social dominance is that subordinates show submissive behaviour and in this way relinquish claims on resources. For this to be favoured by evolution, submissive behaviour should have some benefit. In our model, benefits for subordinates come from uncontested resources (*V*_0_), contested resources (*V*_1_(*k*)) acquired by low-ranking individuals (if there are any), and from situations where high-ranking individuals die, causing ranks to improve for survivors. Even in a seemingly extreme situation, such as *V*_14_ in Fig. 3 (case 13 in Table 2, with *V*_0_ = 0), the group members with low Elo ratings should have some reproductive prospects, even if these are small. Because of mortality during the season, there is a small probability that all or nearly all higher-ranked competitors eliminate each other, leaving a surviving and previously low-ranked individual with reproductive benefits (e.g., in Fig. S2d there are a few data points with positive reproductive success for individuals with low Elo rating). Some chance of reproductive success, even if it is small, can thus select against ‘desperado’ strategies, which otherwise would prevent dominance hierarchies from forming, and might instead promote the evolution of fatal fighting (Enquist and Leimar 1990). For cases with uncontested resources (*V*_0_ > 0), subordinates have greater prospects, which means that submission for low-ranked individuals can be beneficial with little or no mortality for higher ranks.

We have also investigated the consequences of eliminating the risk of death from our model, with the expectation that strategies of refusing to submit should be favoured. From simulations (not shown) we found that without mortality costs (*c*_1_ = 0), dominance hierarchies do not form because individuals keep fighting and reproductive success becomes uncorrelated with fighting ability.

Monopolisation of resources by dominants implies a non-linear relation between rank and reproduction (Fig. 1c). Given that all ranks have some expected reproductive success, the influence of this non-linearity on winner and loser effects can be understood. In our model, winner and loser effects derive from generalisation (see explanation of results in Fig. 4). For instance, for the *V*_14_ cases generalisation is high (Table 2), which makes sense because the lowest-ranked individuals have little to gain from persisting in aggression in order to move up one rank or two, making loser effects adaptive for them. Contrast this with a linear case, where each increase in rank corresponds to the same increment in acquired resources, which holds for the distribution *V*_11_ in Fig. 1c, and also corresponds to the assumptions of the previous model (Leimar 2021). The evolutionary consequence of the linearity is limited generalisation (Table 2) with smaller winner and loser effects, as well as lower costs of fighting (Fig. 3d).

The perspective of social competence (Taborsky and Oliveira 2012; Bshary and Oliveira 2015; Fernald 2017; Varela et al. 2020) is relevant for these winner and loser effects. A key idea is that individuals adjust their fighting behaviour based on the consequences of winning or losing an interaction, using information obtained through learning. The strength of winner-loser effects should then depend on two factors: the relationship between rank and fitness and an individual’s position within the hierarchy. If only top-ranking individuals acquire contested resources, losing once provides information that the top rank is out of reach, which should lead to reduced willingness to fight, in agreement with our results (Fig. 4).

It follows that there are two kinds of fitness non-linearity that are important for the results. First, with mortality as the cost of fighting, costs and benefits are non-additive, and it can be adaptive for low-ranked individuals to be submissive and avoid accumulating damage that would put their reproductive benefits at risk. Second, this effect becomes stronger when there is a non-linear relation between rank and resources.

### Reproductive skew models

Reproductive skew in social groups has been fairly much studied using evolutionary analysis (Johnstone 2000; Port and Kappeler 2010). Of these the ‘tug-of-war’ model (Reeve et al. 1998) shows some similarity to our model here, in that assumptions are made about costs of conflicts in a tug-of-war over reproduction. Even so, there are qualitative differences between our analysis and previous approaches. Our model makes assumptions about uncontested (*V*_0_) and contested (*V*_1_(*k*)) resources, and derives outcomes for reproductive skew and costs of contests from these assumptions, based on the evolution of behavioural mechanisms of hierarchy formation. Importantly, we also assume that the total reproductive output of a local group (e.g., 16 offspring in our simulations) is not influenced by mortality costs of accumulated damage; mortality only affects an individual’s share of the total. For the tug-of-war model, the aim is instead to derive predictions about reproductive skew from assumptions about how costs of conflicts reduce the total reproductive output of a group (which is a pair in the tug-of-war model (Reeve et al. 1998)). Because of these differences in basic assumptions, it is not meaningful to directly compare the results of the models.

Nevertheless, it is of interest to extend our model by incorporating common interest between group members, for instance letting the costs of conflicts reduce the total reproduction by a group. Our current assumption of a constant total reproductive output might be reasonable in some cases, such as for males competing over mating opportunities, but could be less realistic for females competing over resources. It is likely that common interest would lower the cost of hierarchy formation. In addition to common interest, allowing for multi-year life histories is a natural extension, raising the issue of the extent to which interactions from previous years should be remembered. This could throw light on questions of queuing for dominance, for which there are reproductive skew models (Kokko and Johnstone 1999).

Contested and uncontested resources are important ingredients in our model. They represent different assumptions about the distribution of resources in space and time. For instance, the distribution *V*_14_ of contested resources (Fig. 1) could represent competition between males over mating when one or more females become receptive, or the competition between females when there is a single suitable breeding territory. As mentioned, a distribution *V*_1_(*k*) need not directly translate to reproductive skew; if individuals do not submit, acquired resources in a reproductive cycle become random, so with several reproductive cycles, there would be little or no reproductive skew. We might compare with experimental observations of the difficulties for high-ranking male junglefowl to control matings in a group (McDonald et al. 2017), reducing or even eliminating skew. Thus, it is the shape of *V*_1_(*k*) together with a stable hierarchy where subordinates yield to dominants that gives rise to reproductive skew in our model.

The distribution *V*_11_ (Fig. 1c) could represent a situation where resources appear in a dispersed manner, and if they are contested it is typically by two group members. Similarly, *V*_0_ could correspond to resources that individuals encounter singly, or possibly to resources acquired through alternative, non-aggressive strategies. Finally, if nearby territories are distributed according to rank, territory quality will influence the distribution of reproductive success.

### Comparison with observations

There are many simplifying assumptions in our model compared to natural situations. Among these are annual life histories, absence of common interest and relatedness between interacting individuals, no forgetting of previous interactions, and no within-season dispersal to other groups. Even with these simplifications, it is of interest to compare our results with observations, to gain further understanding of the evolution of social dominance.

### Rank and reproductive success

Obtaining data on both lifetime reproductive success and social dominance is challenging, but there are nevertheless many studies. While there is strong support for a general reproductive advantage of higher rank, genetic data also show that monopolisation of mating by dominant males in polygynous mammals is typically not complete (Pemberton et al. 1992; Hogg and Forbes 1997; Coltman et al. 1999; Worthington Wilmer et al. 1999; Coltman et al. 2002; Hoffman et al. 2003; Alberts et al. 2006; Twiss et al. 2006; Wroblewski et al. 2009; Pörschmann et al. 2010; Stopher et al. 2011). It thus seems that the most extreme case among our simulations (case 13 in Table 2, Fig. 1c, Fig. 3) is unrealistic in its assumptions about acquired resources. A certain amount of uncontested resources (*V*_0_) and/or the distributions *V*_11_, *V*_12_ or *V*_13_ might better correspond to natural situations. Combining uncontested resources with monopolisation of contested resources by dominants, we found that the relation between fighting ability and Elo rating was more diffuse for low than for high ranks (Fig. S6d and S7d). This would fit with observations where the precise positions of top-ranking individuals are clear while it is harder to rank individuals lower in the hierarchy.

### Costs of dominance interactions

Our simulations yielded fairly high mortality costs of dominance interactions, also with uncontested resources and limited reproductive skew (Fig. 3d). These results raise the question of whether high costs of hierarchy formation occur in nature. As an example, in female junglefowl, that are similar to the domestic fowl where dominance hierarchies were first described (Schjelderup-Ebbe 1922), a strongly skewed distribution of reproductive success was recorded in a relatively large free-ranging group (Collias et al. 1994), but the costs of hierarchy formation for female junglefowl do not appear to be high. A potential explanation for the seeming discrepancy is that breeding junglefowl occur in rather small groups in nature (Johnson 1963; Collias and Collias 1967), perhaps with less competition over nest sites and other resources. The example illustrates the difficulties of estimating costs of social dominance: these estimates need to come from situations similar to those where the behavioural mechanisms evolved.

The important effect of mortality in our model is that individuals lose further opportunities for reproduction. If individuals instead are driven out of the group and fail to gain reproduction in another group, the evolutionary effect would be the same, and this might be relevant for males in polygynous species. Variation in the duration of tenure of a dominant male in a group is sometimes observed (Hoffman et al. 2003) and could be an indication of costs. The general idea has support from a study of the factors influencing tenure duration in polygynous mammals (Lukas and Clutton-Brock 2014).

Overall, although our results that costs of dominance interactions can be high has potential for agreement with observations, perhaps even more when taking physiological costs into account (Briffa and Sneddon 2007), the issue seems not to be resolved using current data. One thing to note is that our model deals with situations where social hierarchies distribute lifetime reproduction, whereas many observations of dominance interactions come form situations where individuals compete for resources that represent smaller fitness effects.

### Winner and loser effects

We found a dramatic influence of the shape of the distribution of contested resources on winner and loser effects (Fig. 4a, S6a, S7a), and this might ease comparison with observations. Keeping in mind the limitations of our assumptions, our results for non-linear distributions *V*_1_(*k*) could correspond to the behaviour of younger individuals in some species, who quickly learn to avoid contesting higher-ranked opponents, or to individuals pursuing alternative, non-aggressive strategies. Our model would need to be extended to further explore this issue.

There are several experiments on winner and loser effects (Rutte et al. 2006), but they are often on species where the fitness effects of dominance in natural situations are unknown. In addition, the natural breeding situation is frequently that of nearby or partly overlapping territories or sites. To give a few examples, reproductive male three-spined stickleback show stronger loser than winner effects (Bakker et al. 1989), and natural aggressive interactions in this species likely occur between males in nearby territories (Bakker and Sevenster 1983). Experiments with pumpkinseed sunfish have found weak and short-lasting winner effects (Chase et al. 1994) and stronger and more long-lasting loser effects (Beacham and Newman 1987; Beacham 1988). These experiments used fish that were not breeding but taken from mixed-sex shoaling groups of individuals in the field. The social structure of breeding pumpkinseed sunfish is likely to be colonies of male nests that females visit and lay eggs in. Finally, there are several studies on aggression and social dominance in the green swordtail, including on bystander, winner, and loser effects (Earley and Dugatkin 2002), as well as field studies on reproductive skew (Tatarenkov et al. 2008). The social system of the species in nature is partially overlapping male home ranges, sometimes with the presence of lower-ranking ‘satellite’ males, somewhat similar to leks (Franck and Ribowski 1993; Franck et al. 1998). These studies illustrate that it could be challenging to link experimental work on social dominance to field situations, including to fitness effects of rank. There is broad agreement between the studies and results from our current (Fig. 4) and previous (Leimar 2021) models in that loser effects are typically stronger than winner effects, but the differences between experiments and field situations make it harder to evaluate whether effects of non-linearity of the distribution of resources over ranks, like those in Fig. 4, occur in nature.

### Consistent behaviour

Traits related to aggression can be viewed as components of animal personalities (Sih et al. 2004; Réale et al. 2007), which are characterised by consistency over time and contexts. For situations with pronounced monopolisation of contested resources by dominants, like the *V*_12_ and *V*_14_ distributions (Fig. 1c), our analysis predicts substantial individual differences in aggressiveness, as a consequence of generalisation from learning in a group (Fig. 4, S6, S7, Table 2). Distribution like *V*_11_ or *V*_13_ would instead result in weaker such effects. The importance of social experience, compared to genetic or developmental variation, is of interest to research on animal personalities, and our results can provide some insight. Still, just as for winner and loser effects, there is the difficulty of linking observations to fitness effects in the field, in particular to the shape of distributions of contested resources over ranks, limiting the conclusions that can be drawn.

In male junglefowl, winner and loser effects appear to have only a weak influence on dominance hierarchy formation, whereas genetic or developmental variation in aggressiveness is more important and shows consistency over time (Favati et al. 2017, 2021; Pizzari and McDonald 2019). These observations would agree with our results, provided that the distribution of acquired resources has roughly a linear shape. A linear shape is consistent with experimental data (McDonald et al. 2017), but it is not known how well these results correspond to field situations. Similar conclusions could apply to certain swordtail species (Wilson et al. 2011, 2013; Boulton et al. 2018). It would be of interest to investigate species with pronounced monopolisation of contested resources, in order to test whether they show strong effects of social experience on aggressiveness.

### Modelling social dominance and aggression

The game theory used here investigates the evolution of traits that control behavioural mechanisms, such as the parameters of actor-critic learning, over a range of situations. In terms of social cognition, the approach is in-between behaviourist assumptions of universal cognitive mechanisms and those of traditional game-theory modelling, entailing that individuals make optimal decisions in each particular situation. The overall aim of the approach is to integrate function and mechanism (McNamara and Houston 2009).

The issue of modelling styles can be formulated as a distinction between small-worlds models, in which individuals have accurate innate representations of their environment, including the decision-making machinery of others, and large-worlds models, where individuals instead rely on limited cognitive mechanisms that nevertheless have proven their worth over a range of imperfectly delineated circumstances (McNamara and Leimar 2020). There is also a correspondence to ideas about social competence (Taborsky and Oliveira 2012; Bshary and Oliveira 2015; Fernald 2017), focusing on how general developmental and cognitive mechanisms become adapted to social situations, thereby allowing individuals to respond effectively to their environment.

There is a long tradition of using learning in game theory, both in biology (Harley 1981) and economics (Fudenberg and Levine 1998). This can be combined with a study of the evolution of learning parameters or traits (Niv et al. 2002; Hamblin and Giraldeau 2009; Dridi and Lehmann 2014), as we have done here. To promote realism, it is preferable to study learning traits that have a correspondence in animal psychology and neuroscience. For instance, aggressive behaviour is regarded as a potentially rewarding activity in animal psychology (Hogan and Roper 1978; Domjan et al. 2000; Fish et al. 2005). We implemented this perspective through a perceived reward *v_i_* of performing aggression (Table 1). In traditional small-worlds game theory, rewards and payoffs are treated as the same and payoffs have the interpretation of fitness increments. In our model, as well as in similar learning models (McNamara and Leimar 2020; Leimar 2021; McNamara et al. 2021), perceived rewards and penalties of aggressive behaviour are instead interpreted as components of a mechanism that functions to guide an individual’s life history in a favourable manner. Thus, the simulations in Table 2 with *V*_0_ = 9 have smaller evolved values of *v*_i_ than those with *V*_0_ = 0,1, 4, and also lower preference learning rates *α_θi_* and initial aggression parameters *θ*_0*i*_, all of which tend to make individuals act in a cautious manner, reducing the risk of losing their substantial uncontested resources.

Our current model could be extended to include elements like bystander observations (Leimar 2021), multi-year life histories, dispersal, territoriality, or relatedness between group members. Among the ingredients needed for this to succeed are reasonable specifications of traits and perceptions of the interacting individuals. We believe such endeavours benefit from collaboration between modellers, experimentalists, and biologists with experience from the field, because this helps overcoming the considerable challenges of linking theoretical constructs to natural situations.

## Competing interests

The authors declare no competing interests.

## Funding

This work was supported by a grant (2018-03772) from the Swedish Research Council to OL.

## Supplementary information

### Additional figures

**Figure S1:**
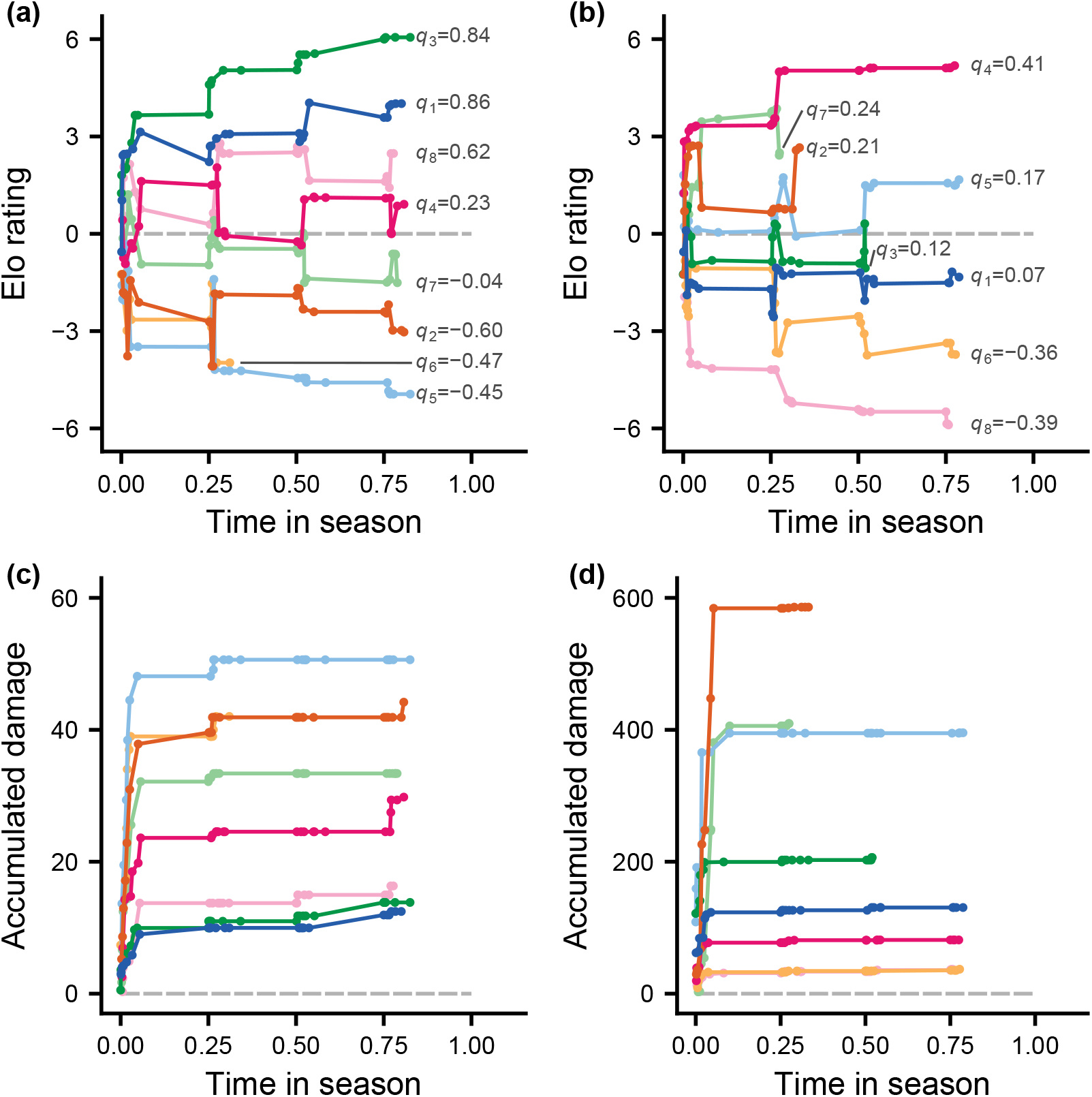
Panels (a) and (c) show the Elo rating and the accumulated damage over the mating season for one example of a group of 8 individuals, with AR parameters as for case 1 in Table 2 and learning traits given by the mean values for this case. The curves have different colours to allow comparison between panels (a) and (c), and are labelled in (a) with the fighting ability *q_i_* of the individual. Panels (b) and (d) show the same for case 13 in Table 2.

**Figure S2:**
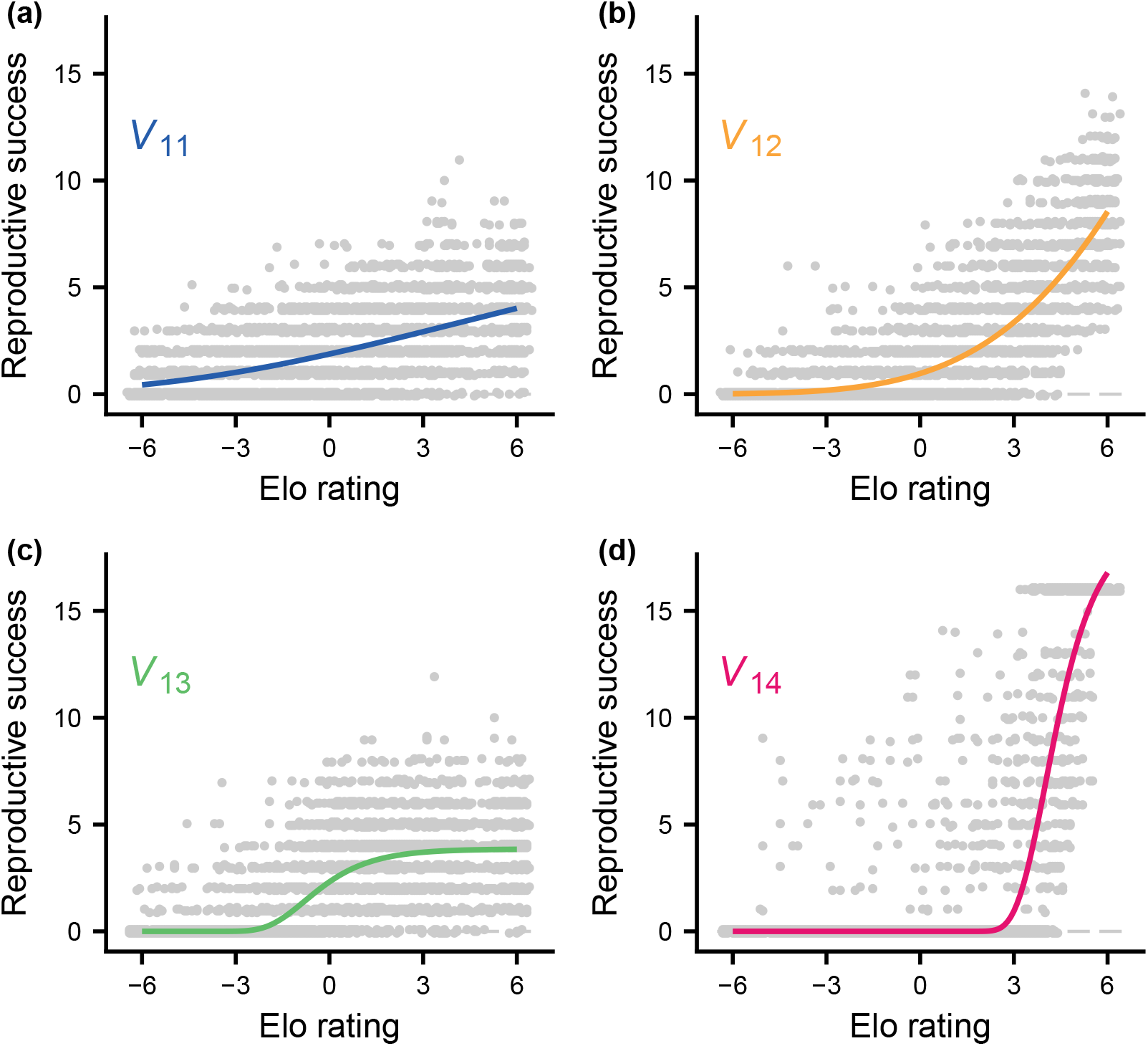
Simulated data and fitted curves of RS (number of offspring) for the cases in Fig. 3a (colour coded). Panels (a) to (d) show cases 1, 5, 9, and 13 in Table 2. Non-linear regression (nls function in R) was used to fit RS as a Gompertz function of the Elo rating, as described below. The locations of the grey points are shifted by small random amounts to reduce overlap.

**Figure S3:**
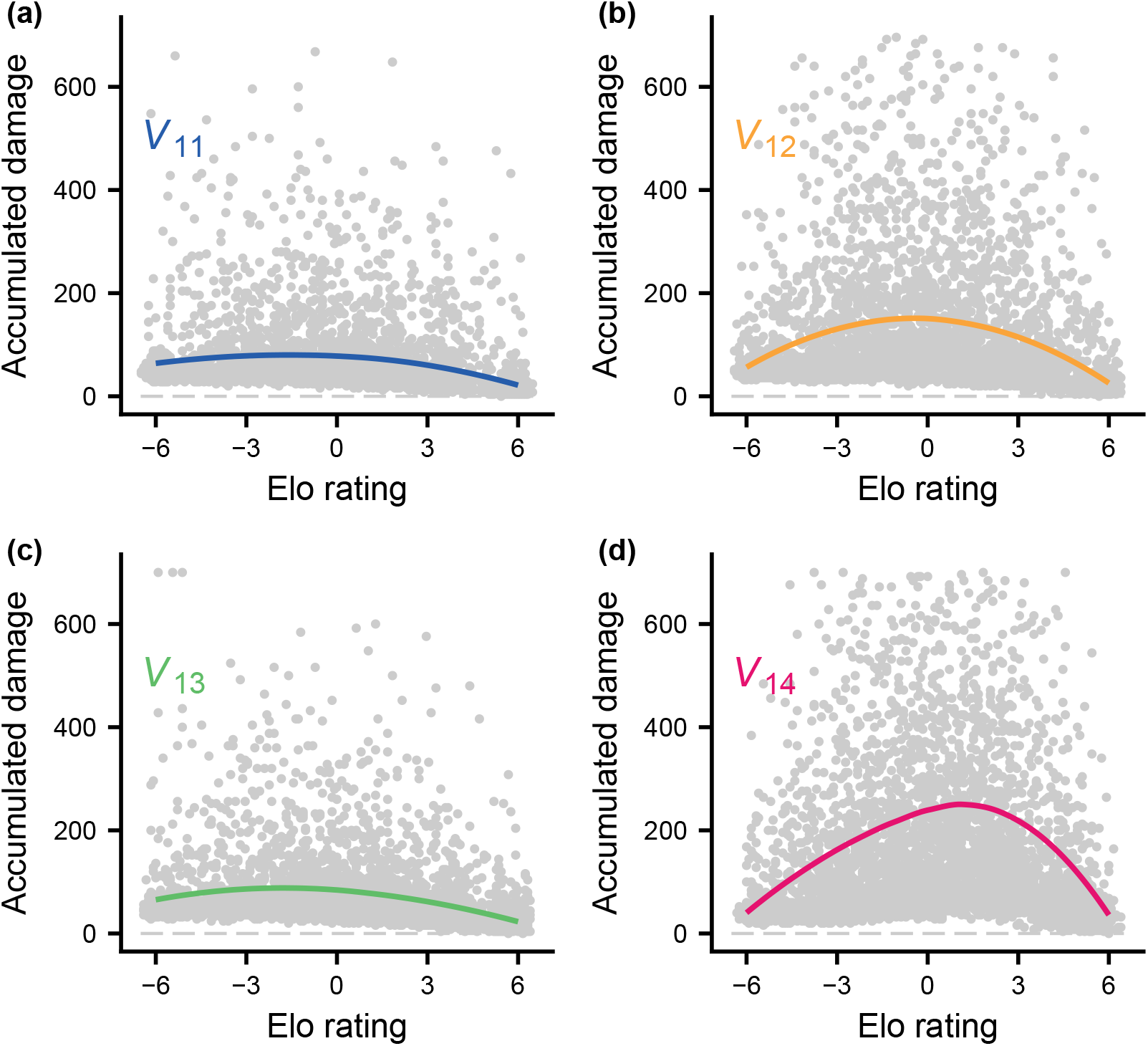
Simulated data and fitted curves of accumulated damage over the mating season for the cases in Fig. 3c (colour coded). Panels (a) to (d) show cases 1, 5, 9, and 13 in Table 2. Local regression (loess function in R) was used to fit accumulated damage *D_a_* as a function of the Elo rating, as described below. The locations of the grey points are shifted by small random amounts to reduce overlap.

**Figure S4:**
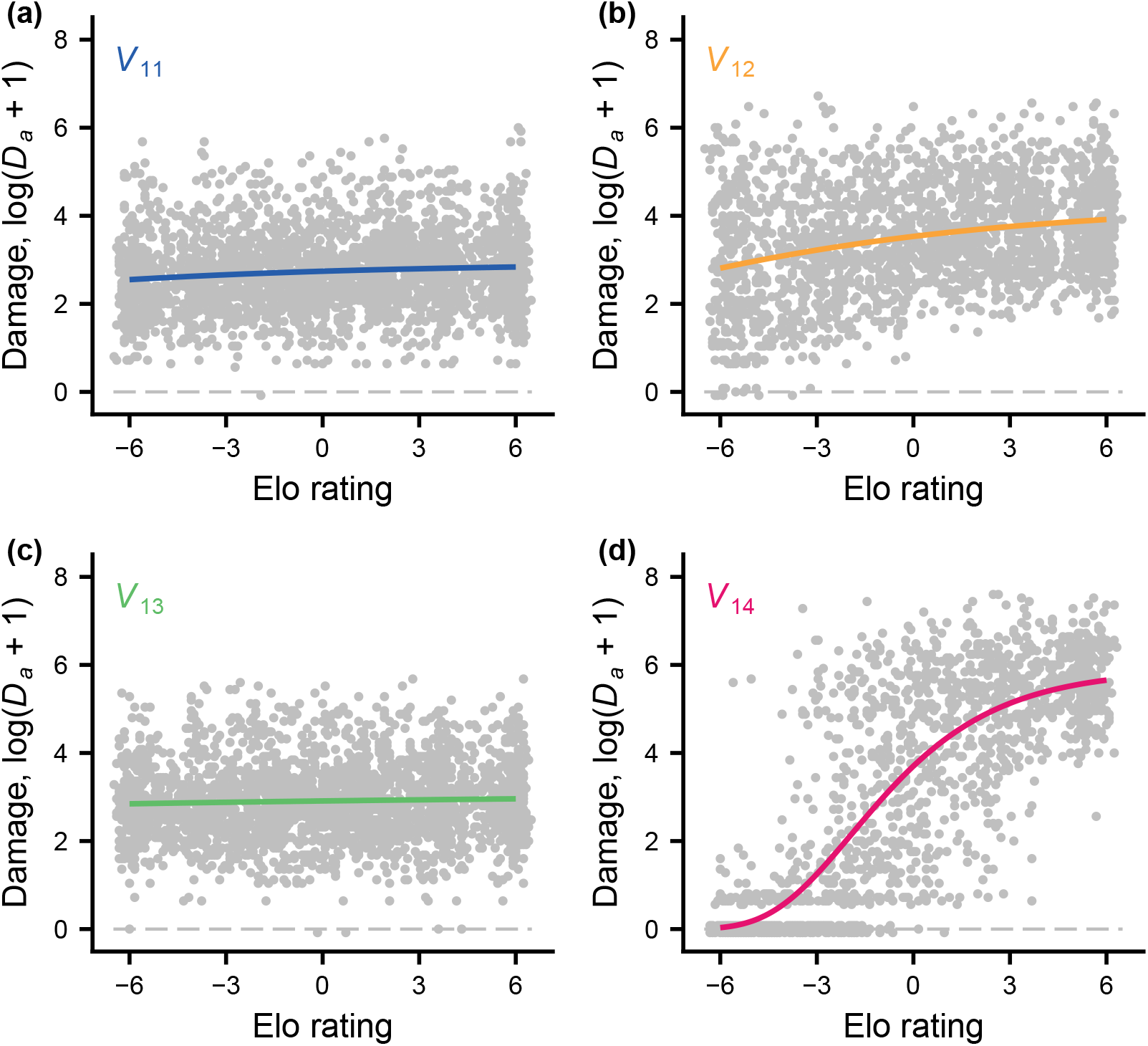
Simulated data and fitted curves of accumulated damage in staged winner-loser experiments against new, matched individuals for the cases in Fig. 4b (colour coded). Panels (a) to (d) show cases 1, 5, 9, and 13 in Table 2. Non-linear regression (nls function in R) was used to fit log(*D_a_* + 1) as a Gompertz function of the Elo rating, as described below. The locations of the grey points are shifted by small random amounts to reduce overlap.

**Figure S5:**
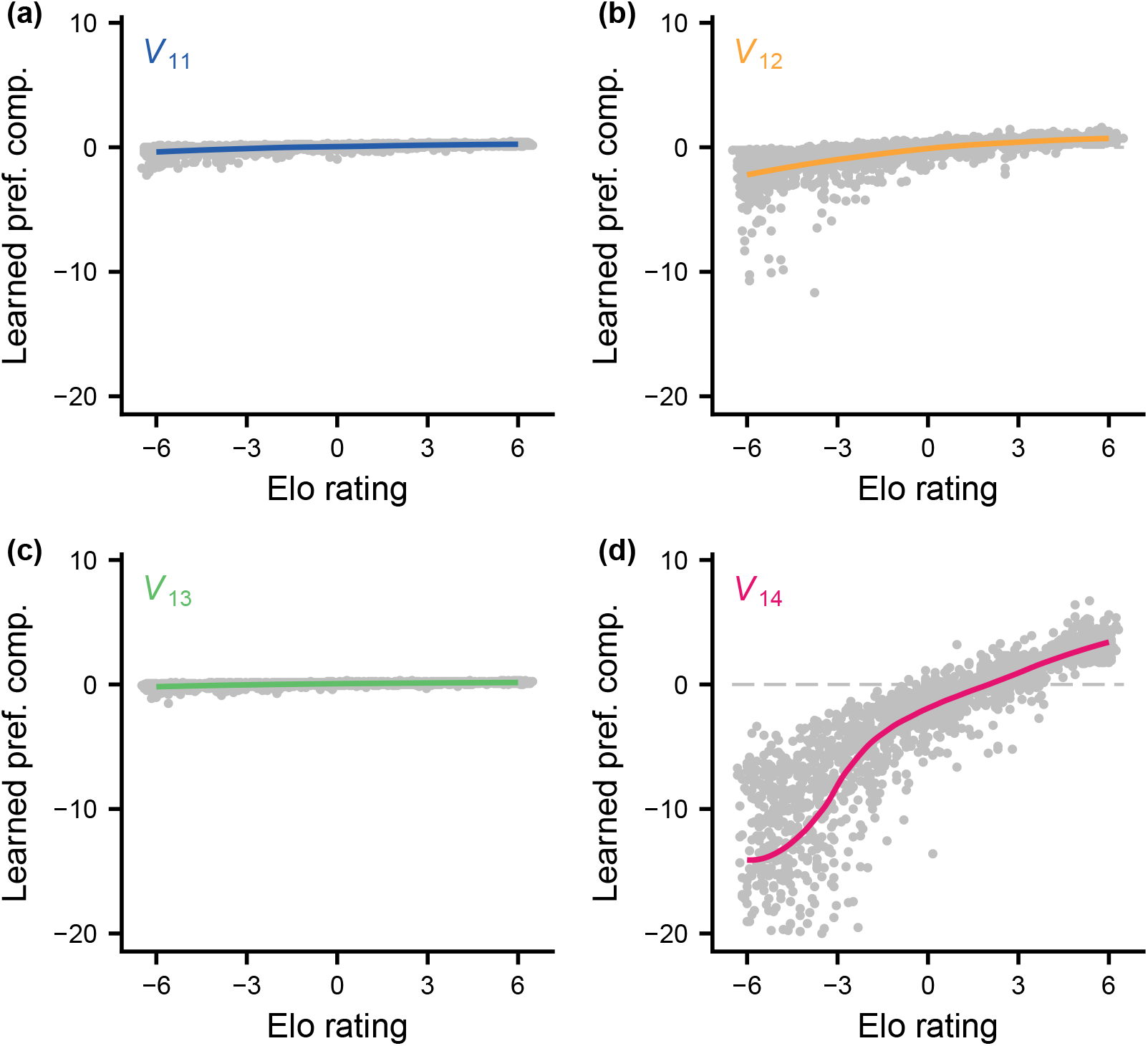
Simulated data and fitted curves of the generalised preference component (*h_iit_* in Table 1) at the end of the mating season for the cases in Fig. 4c (colour coded). Panels (a) to (d) show cases 1, 5, 9, and 13 in Table 2. Local regression (loess function in R) was used to fit the generalised preference component as a function of a group member’s Elo rating at the end the mating season. The locations of the grey points are shifted by small random amounts to reduce overlap.

**Figure S6:**
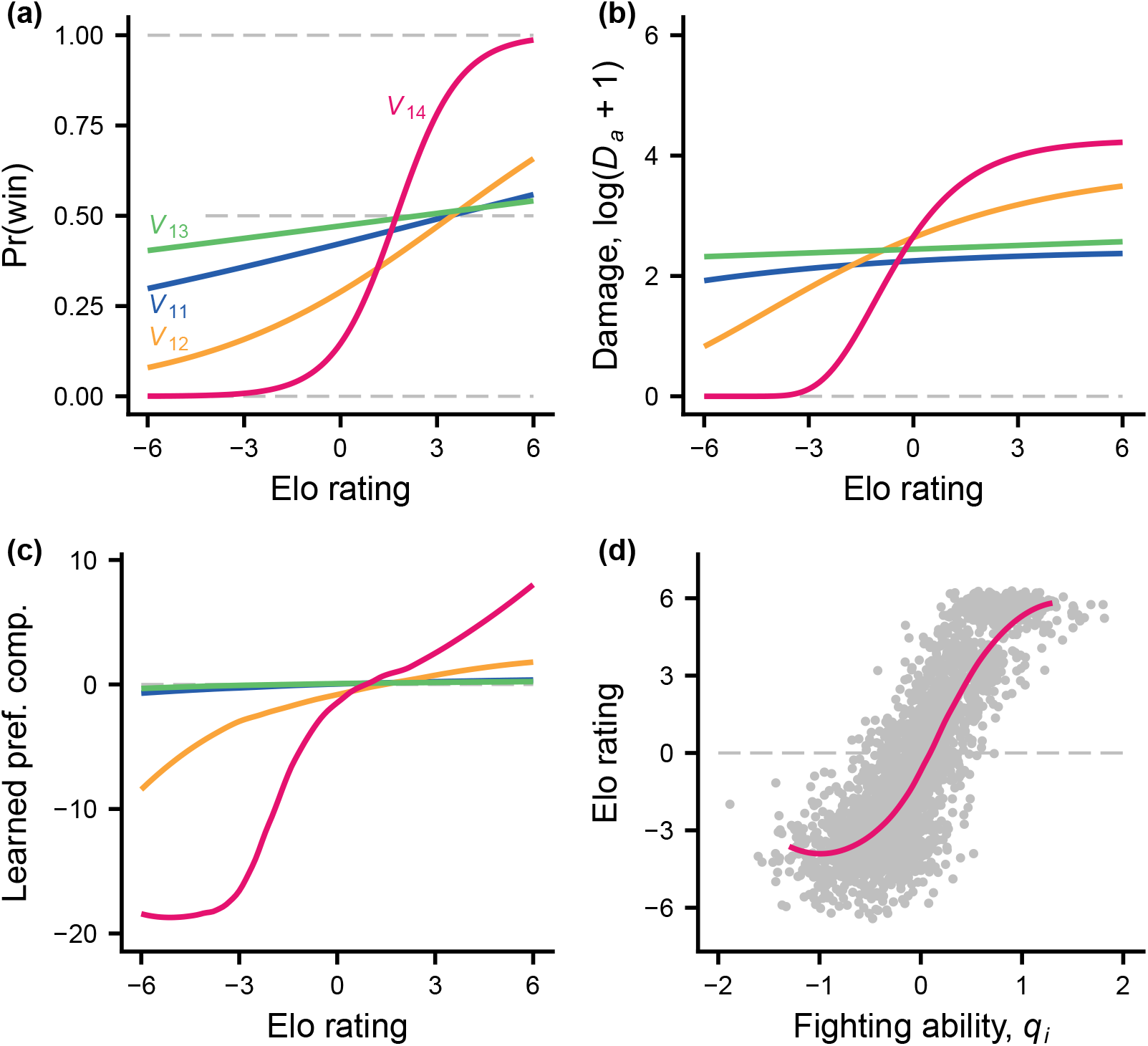
Illustration of hypothetical winner- and loser-effect experiments, as in Fig. 4, but with uncontested AR. Each group member that survived over the mating season had a staged interaction with a matched (equal fighting ability, *q_i_* = *q_j_*) new and naive opponent. A staged pair had up to 10 contests as described in Fig. 1b, ending when dominance was settled. Cases 2, 6, 10, and 14 in Table 2 are shown (colour coded after the shape of *V*_1_), all of which have *V*_0_ = 1. For each case there were 500 simulated groups, including winner-loser experiments. (a) Fitted (logistic regression) probability of winning (becoming dominant) for a group member interacting with a matched, naive opponent, as a function of the group member’s Elo rating. (b) Fitted accumulated damage (logarithmic scale) from these contests. (c) Fitted generalised action preference component (*h_iit_* in Table 1) for group members at the start of staged interactions, as a function of their Elo rating. (d) Data and a loess fitted curve of the Elo rating as a function of fighting ability *q_i_* of group members, for case 14 in Table 2 (i.e., *V*_14_ in panels a-c).

**Figure S7:**
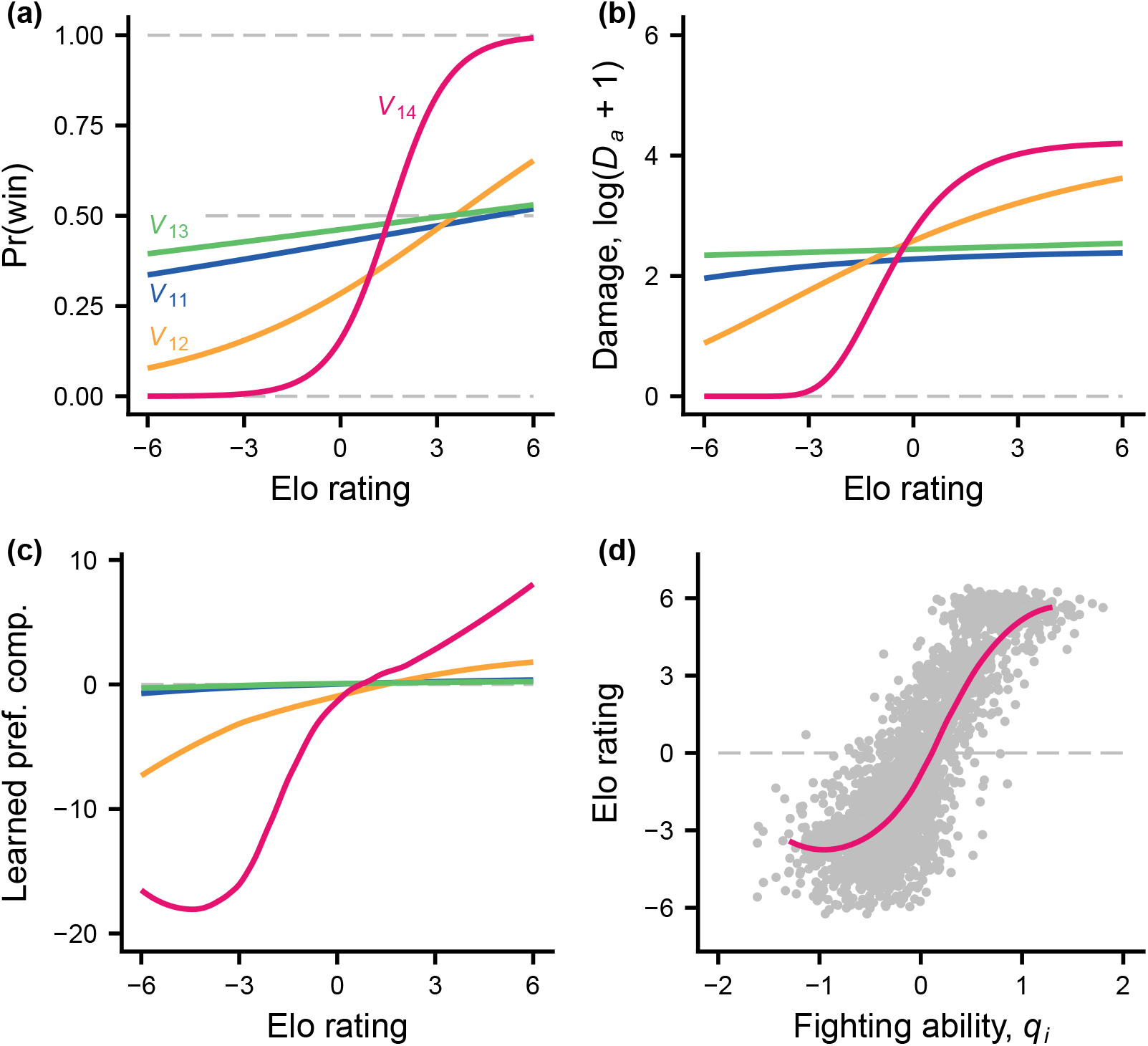
Illustration of hypothetical winner- and loser-effect experiments, as in Fig. 4, but with uncontested AR. Each group member that survived over the mating season had a staged interaction with a matched (equal fighting ability, *q_i_* = *q_j_*) new and naive opponent. A staged pair had up to 10 contests as described in Fig. 1b, ending when dominance was settled. Cases 3, 7, 11, and 15 in Table 2 are shown (colour coded after the shape of *V*_1_), all of which have *V*_0_ = 4. For each case there were 500 simulated groups, including winner-loser experiments. (a) Fitted (logistic regression) probability of winning (becoming dominant) for a group member interacting with a matched, naive opponent, as a function of the group member’s Elo rating. (b) Fitted accumulated damage (logarithmic scale) from these contests. (c) Fitted generalised action preference component (*h_iit_* in Table 1) for group members at the start of staged interactions, as a function of their Elo rating. (d) Data and a loess fitted curve of the Elo rating as a function of fighting ability *q_i_* of group members, for case 15 in Table 2 (i.e., *V*_14_ in panels a-c).

### Statistical fitting of curves to simulated data

Fitted curves are shown in Figs. 2 to 5, and S2 to S7. Non-linear regressions from the R statistical package (version 4.0.5; https://www.R-project.org/) were used, with the aim of giving a reasonable visual summary of the individual data points from simulations. The variation around the fitted curves can be judged from Figs. S2 to S5.

We used either the nls function, (Figs. 3a, 4b, 5a, 5b, S2, S4, S6b, S7b), the `loess` function (Figs. 3c, 4c, 4d, 5c, 5d, S3, S5, S6c, S6d, S7c, S7d), or logistic regression using the `glm` function (Figs. 4a, S6a, S7a). For the nls fits of RS to Elo rating *E_i_* we used either a Gompertz function, *A* exp(—*B* exp(−*CE_i_*)), or a modified Gompertz function *A* exp (—*B* exp(—*CE_i_*) + *D*, as the non-linear function, with *A,B,C,D* as parameters to be fitted.

### Model details

Several parts of the model are the same as in a previous one (Leimar 2021). These are the observations in a round, the actions A and S, the action preferences and estimated values, the implementation of action exploration, and the learning updates. They are described previously (Leimar 2021) but, for completeness, they are also given here. Aspects of the current model that are different from the previous one are outlined in the main text, including in Table 1, and at the end of this section we give additional details.

### Observations and actions

The model simplifies a round of interaction into two stages. In the first stage, interacting individuals make an observation. Thus, individuals observe some aspect *ξ* of relative fighting ability and also observe the opponent’s identity. The observation by an individual is statistically related to the difference in fighting ability between itself and the opponent, *q_i_* — *q_j_*. For the interaction between individuals *i* and *j* at time *t*, the observation is

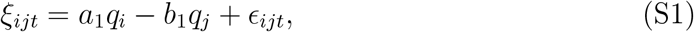

where *a*_1_,*b*_1_ ≥ 0 and *ϵ_ijt_* is an error of observation, assumed to be normal with mean zero and SD *σ* (*a*_1_ = *b*_1_ is assumed in all simulations). By adjusting the parameters *σ_q_*, which is the SD of the distribution of *q_i_*, and *a*_1_, *b*_1_, and *σ* from equation (S1), one can make the information about relative quality more or less accurate. The observation (*ξ_ij_, j*) is followed by a second stage, where individual *i* chooses an action, and similarly for individual *j*. The model simplifies to only two actions, A and S, corresponding to aggressive and submissive behaviour.

### Action preferences and estimated values

For an individual *i* interacting with *j* at time *t, l_ijt_* denotes the preference for A. The probability that *i* uses A is then

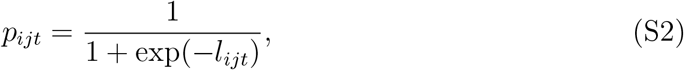

so that the preference *l_ijt_* is the logit of the probability of using A. The model uses a linear (intercept and slope) representation of the effect of *ξ_ijt_* on the preference, and expresses *l_ijt_* as the sum of three components

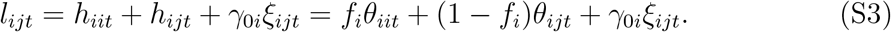

Here *h_iit_* = *f_i_θ_iit_* is a contribution from generalisation of learning from all interactions, *h_ijt_* = (1 — *f_i_*)*θ_ijt_* is a contribution specifically from learning from interactions with a particular opponent *j*, and *γ*_0*i*_*ξ_ijt_* is a contribution from the current observation of relative fighting ability. Note that for *f_i_* = 0 the learning about each opponent is a separate thing, with no generalisation between opponents, and for *f_i_* = 1 the intercept component of the action preference is the same for all opponents, so that effectively there is no individual recognition (although the observations *ξ_ijt_* could still differ between opponents). One can similarly write the estimated value 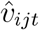 of an interaction as a sum of three components:

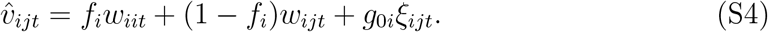

The actor-critic method updates *θ_iit_, θ_ijt_, w_iit_*, and *w_ijt_* in these expressions based on perceived rewards, whereas *f_i_, γ*_0*i*_, and *g*_0*i*_ are genetically determined.

### Exploration in learning

For learning to be efficient over longer time spans there must be exploration (variation in actions), in order to discover beneficial actions. Learning algorithms, including the actor-critic method, might not provide sufficient exploration, because learning tends to respond to short-term rewards. In the model, exploration is implemented as follows: if the probability in equation (S2) is less than 0.01 or greater than 0.99, the actual choice probability is assumed to stay within these limits, i.e. is 0.01 or 0.99, respectively. In principle the degree of exploration could be genetically determined and evolve to an optimum value, but for simplicity this is not implemented in the model.

### Perceived rewards

An SS interaction is assumed to have zero rewards, *R_ijt_* = *R_jit_* = 0. For an AS interaction, the aggressive individual *i* perceives a reward *R_ijt_* = *v_i_*, which is genetically determined and can evolve. The perceived reward for the submissive individual *j* is zero, *R_jit_* = 0, and vice versa for SA interactions. If both individuals use A, some form of costly dominance display or fight occurs, with perceived costs (negative rewards or penalties) that are influenced by the fighting abilities of the two individuals. The perceived rewards of an AA interaction are assumed to be

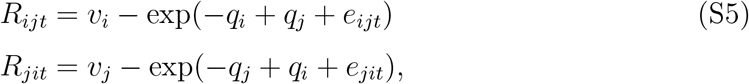

where *e_ijt_* is a normally distributed random influence on the perceived penalty, with mean zero and standard deviation *σ_p_*, and similarly for *e_jit_*.

### Learning updates

In actor-critic learning, an individual updates its learning parameters based on the prediction error (TD error)

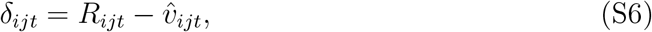

which is the difference between the actual perceived reward *R_ijt_* and the estimated value 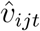. The learning updates for the *θ* parameters are given by

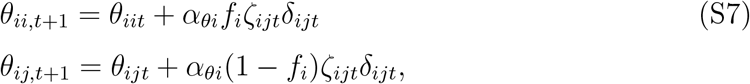

where

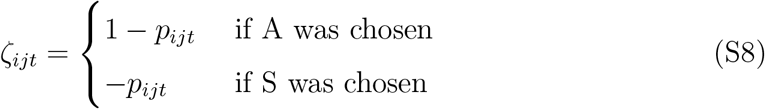

is referred to as a policy-gradient factor and *α_θi_* is the preference learning rate for individual *i*. Note that *ζ_ijt_* will be small if *p_ijt_* is close to one and individual *i* performed action A, which slows down learning, with a corresponding slowing down if *p_ijt_* is close to zero and S is chosen. There are also learning updates for the *w* parameters given by

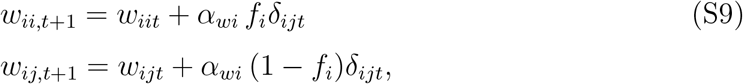

where *α_wi_* is the value learning rate for individual *i*.

The updates to the policy parameters *θ* can be described using derivatives of the logarithm of the probability of choosing an action with respect to the parameters. Using equation (S2), we obtain

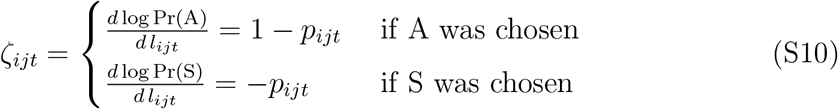

for the derivative of the logarithm of the probability of choosing an action, A or S, with respect to the preference for A, which corresponds to equation (S8). From equation (S3) it follows that

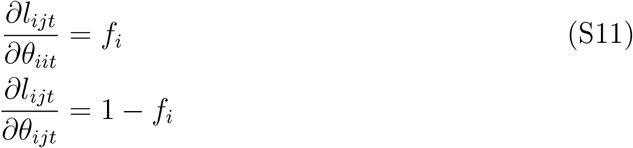

and this gives the learning updates of the *θ* parameters in equation (S7). The updates of the *w* parameters of the value function can also be described using derivatives. From equation (S4) it follows that

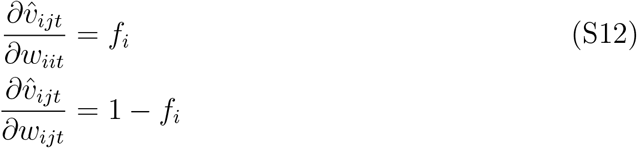

and this gives the learning updates of the *w* parameters in equation (S9).

### Life-history and reproductive season

There is an annual life cycle with a single reproductive season. Dominance interactions occur in groups of size *g_s_*, with *g_s_* = 8 for the individual-based simulations in Table 2. The season is split into a number of reproductive cycles; there are 4 cycles for the simulations in Table 2. A cycle starts with a sequence of contests. Each contest is between a randomly selected pair of group members and there are 5*g_s_*(*g_s_* — 1) contests in a cycle, i.e., on average 10 contests per pair. As a result of the contests, a dominance hierarchy is formed, and group members acquire resources according to their ranks. Survival from one cycle to the next depends on the damage accumulated in contests. The purpose of the scheme is to implement a combination of hierarchy formation, resource acquisition, and mortality over the season in a way that allows both fitness benefits and costs to influence trait evolution. In principle very similar results could be achieved by, for instance, implementing a risk of mortality after each contest, or even after each round of interaction.

### Contests

If a dominance relation has already been established between contestants *i* and *j* in the current cycle, there is no interaction. If not, the contestants go through a number of rounds, at minimum 10 rounds and at maximum 200 rounds of interaction. If there are 5 successive rounds where *i* uses A and *j* uses S (5 AS rounds), the contest ends and *i* is considered dominant over *j*, and vice versa if there are 5 successive SA rounds. Further, the contest ends in a draw if there are 5 successive SS rounds. In the following cycle (if there is one), previously dominance relations are reset, so the hierarchy needs be reformed, but group members retain their learning. The reason for this is to induce a kind of exploration of the dominance relations. Previously undecided relations can then become decided, and the possible consequences of group members dropping out (dying) between one cycle and the next can have an influence.

### Acquired resources

After the contests in the current cycle, group members acquire resources. Each (surviving) individual obtains an increment *V*_0_, irrespective of dominance relations. If a linear hierarchy has been established, an individual with rank *k* (with *k* =1 the top rank) obtains a resource increment of *V*_1_(*k*). The ranking is based of how many other group members an individual dominates (this measure is referred to as a score structure by Landau (1951)). If some individuals dominate the same number of other group members, their relative rank is randomly determined. So, for instance, if all individuals would use action S in the contests, there would be no real dominance hierarchy (each would dominate 0 other group members), and it is random which of them obtains *V*_1_(1), *V*_1_(2), etc.

### Accumulated fighting damage and mortality

A group member *i* accumulates damage *D_ait_* from fighting. In an AA round between *i* and *j*, the increment to *D_ait_* is

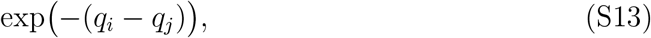

and similarly for *j*. An individual with accumulated damage *D_ait_* survives from one cycle to the next with probability

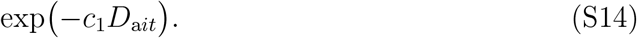

### Reproductive success

A local group, containing 8 interacting individuals and 8 of the other sex, produces 16 offspring. For each offspring, one parent of each sex is randomly drawn from the group, with a probability proportional to acquired resources for interacting individuals. In the next generation, each offspring disperses to a random local group. In this way, interacting individuals are unrelated.

### Elo rating

Several approaches to Elo ratings have been used, differing in such things as the zero point of the scale and the amount to change ratings after a ‘win’ by one individual over another, or after a ‘draw’. There is similarity between updates of Elo ratings and the updates of action preferences for actor-critic learning described above. Here, however, we use the Elo rating just as a conventional measure or index of dominance rank, without further interpretation of what the scores might mean. The possible usefulness of this measure needs instead to be investigated. From our results here, Elo ratings appear useful in providing a simplified description of a dominance hierarchy.

Let *E_it_* be the Elo rating of group member *i* at time *t*. Initially all have rating *E*_*i*0_ = 0. If a contest between *i* and *j* ends with *i* becoming dominant over *j, E_it_* is incremented by

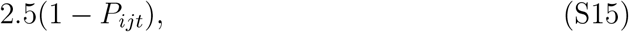

where

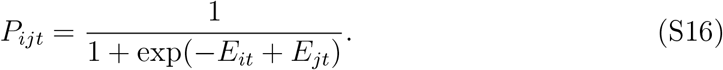

The Elo rating of *j, E_jt_*, is decremented by the same amount. If the contest ends in a draw, *E_it_* is decremented by

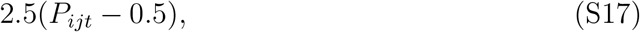

and *E_jt_* is incremented by this amount. It can help the interpretation to think of *P_ijt_* as the probability, before the interaction, of the outcome (‘win’, ‘loss’, or ‘draw’). This, however, is just an interpretation that helps explaining why Elo ratings are defined in a certain way. For dominance relations, which are qualitatively different from wins and losses in a tournament, it is not certain that Elo ratings are useful for predicting outcomes of dominance interactions. One can, of course, investigate the usefulness for each particular case.

## Notes

### Competing Interest Statement

The authors have declared no competing interest.

